# Mechanical causes and implications of repetitive DNA motifs

**DOI:** 10.1101/2024.04.06.588425

**Authors:** Paul Torrillo, David Swigon

## Abstract

Experimental research suggests that local patterns in DNA sequences can result in stiffer or more curved structures, potentially impacting chromatin formation, transcription regulation, and other processes. However, the effect of sequence variation on DNA geometry and mechanics remains relatively underexplored. Using rigid base pair models to aid rapid computation, we investigated the sample space of 100 bp DNA sequences to identify mechanical extrema based on metrics such as static persistence length, global bend, or angular deviation. Our results show that repetitive DNA motifs are overrepresented in these extrema. We identified specific extremal motifs and demonstrated that their geometric and mechanical properties significantly differ from standard DNA through hierarchical clustering. We provide a mathematical argument supporting the presence of DNA repeats in extremizing sequences. Finally, we find that repetitive DNA motifs with extreme mechanical properties are prevalent in genetic databases and hypothesize that their unique mechanical properties could contribute to this abundance.

## Introduction

Approximately 70% of the human genome is transcribed [1], leaving at least 25% of the human genome as noncoding DNA, with this proportion potentially being higher if transcriptional noise is considered. Noncoding DNA plays a variety of roles, including serving as regulatory elements such as promoters, enhancers, and silencers. Some regulatory elements function through mechanical deformation, as exemplified by repression via looping in the lac operon [2][3] and the L-arabinose BAD operon [4]. Additionally, in eukaryotes, DNA may regulate transcription by influencing chromatin formation. For instance, certain repeated sequences exhibit differential nucleosome binding abilities, which can have consequences in human disease [5]. Given that these regulatory actions necessitate physical deformation of DNA, further investigation into the relationship between DNA sequence composition and mechanical properties is warranted.

Sequence-dependent properties of DNA are well-established [6], and studies have proposed the concept of a "mechanical code" to describe their significance [7]. For example, strands with high GC content have greater thermal stability due to more favorable stacking interactions of GC base pairs compared to AT base pairs [8]. This property has been suggested to correlate with an increase in persistence length as GC content increases [9]. It is natural to question whether increasing AT content has the opposite effect on persistence length. However, conflicting reports have emerged regarding the flexibility of poly AT sequences, with some studies suggesting that they repel nucleosomes [10] while others indicate that they can easily loop [11]. This discrepancy could arise because certain AT-rich motifs are highly flexible, while others are relatively inflexible. For instance, poly A sequences have been found to be very stiff [12]. Another related topic is fragile sites [13], which often involve repetitive nucleotide motifs.

The formation of hairpin loop secondary structures through complementary base pairing is thought to contribute to the fragility of these sites. Although the ability to form complementary base pairs is a chemical property, the efficiency of hairpin loop formation is likely influenced by the mechanical properties of the sequence. Fragile sites may play a significant role in human biology, as they can rapidly acquire mutations, potentially contributing to oncogenesis. However, in bacteria, these sites might confer a selective advantage via phase switching [14].

Another well-known example of sequence affecting function is observed in CpG steps. A notable feature of CpG steps is their significant depletion in vertebrate genomes, such as the human genome [15]. This depletion is likely due to the propensity of these base pairs to undergo methylation and subsequent deamination [16]. Sequences rich in CpG steps (CpG islands) tend to be exceptionally rigid [6], though CpG steps themselves are more variable in terms of flexibility [17].

Finally, A-tracts represent a particularly significant example of sequence influencing structure in the context of our work. Since the 1980s, A-tracts have been a subject of interest due to their ability to contribute to intrinsic DNA bending, especially when in phase with DNA’s natural periodic helicity [18][19]. As A-tracts are intrinsically bent and DNA regulation relies on bending and looping, they are natural candidates for regulatory sequences [20]. However, the definition of A-tracts is relatively vague, often described as "DNA sequences consisting of four or more consecutive A:T base pairs without a TA step" [21], which remains a common and current definition. This raises the question of whether a more specific definition of A-tracts could provide a sequence with greater curvature or if other sequence motifs might exhibit even more pronounced bending.

With these questions in mind, we conducted a computational survey of the DNA sequence space, aiming to identify structural or deformational extrema. By discovering extrema, such as the most bent and straightest sequences, we can corroborate current findings, more rigorously define sequences with unique mechanical properties, and potentially uncover new sequences of mechanical interest. To obtain data on DNA structural and mechanical properties, we initially relied on two fundamentally distinct studies: one based on experimental observations of X-ray crystal structures of DNA [22], and the other on molecular dynamics simulations of all-atom DNA structures [23]. While this work was in progress, some newer DNA models were released that were computationally efficient and attempted to account for nonlocal effects. A particularly useful methodology was cgDNA+ [24], which we incorporated in analysis with the molecular dynamics and X-ray data.

In pursuing this goal, we discovered evidence corroborating that sequences such as poly A and A-tracts, as well as other repetitive sequences, are indeed mechanical and/or structural extrema. Interestingly, the prevalence of repetitive sequences in biology remains an open question, leading to their occasional designation as the dark matter of DNA [25]. We propose the hypothesis that the unique mechanical properties of repetitive DNA could contribute to their prevalence.

To support this conclusion, we demonstrate that a search of DNA sequence space reveals repetitive sequences as extrema, and these extrema are significantly structurally different from non-extrema. Under certain assumptions, it is possible to mathematically prove that extrema should consist of repetitive motifs. Furthermore, we show that varying repeated motifs result in different levels of prevalence in the BLAST database, with motifs oriented towards mechanical extrema appearing more frequently.

## Methods

### DNA Model

Theoretical studies of DNA dynamics have been traditionally based on worm-like chain approximation of polymer dynamics [26], but this approach does not account for sequence. The first sequence dependent DNA mechanical models [27][22] were based on the Cambridge University Engineering Department Helix Computation Scheme [28]. Modern approaches employ molecular dynamics simulations, and sequence effects on mechanical properties have been investigated using these techniques [12]. Coarse graining is often used to obtain results more quickly [29].

Since all-atom molecular dynamics systems are computationally intensive, they are typically used to study the configuration of sequences of length less than 30 bp with possible small variations. In this project, however, the goal is to search the full sequence space of a 100bp DNA segment, which includes over 8 × 10^59^ elements. We have therefore chosen to utilize the quickest possible method for predicting structure from sequence: a rigid-base pair model [28]. We initially used the rigid-base pair model with two sets of parameters, one from X-ray crystallography [22] and one from molecular dynamics [23].

However, there are naturally drawbacks to utilizing a rigid base pair model with predefined parameters as it is known that the structure of DNA sequences is determined by more than just single base pair steps because of nearest neighbor interactions or base pair deformations because of nonlocal effects like frustration [30][31]. Thus, we included the non-local cgDNA+ model into our workflow [24]. It is slightly slower than the models without nonlocal interaction (5-10x slowdown) so we chose to adjust the search algorithms accordingly. We chose to mostly focus on and compare the results from cgDNA+ and the crystal parametrization in the main text, as the molecular dynamics based parameters have been shown to have limitations in accuracy [32]. However, these results do serve to check to the extent parametrization effects results so we do include a short summary of them in the results and found sequences are in the supplementary information.

### DNA structure descriptors

To identify sequences of mechanical interest, we utilized a variety of metrics, most of which attempt to quantify the intrinsic stiffness and bending of DNA. The most pertinent metric among these is the persistence length, as described below.

*Persistence length* quantifies the deformation of a polymer by measuring the decay of segment correlation as length increases. For DNA, it is common to describe persistence length as either static (*T_s_*), dynamic (*T_d_*), or apparent (*T_p_*) [33][12]. *Static persistence length* is determined on the basis of intrinsic structure without thermal fluctuations, while apparent persistent length takes both intrinsic structure and thermal fluctuations into account. *Dynamic persistence length* can then be recovered from the equation

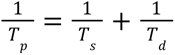

The standard equation for static and apparent persistence length can be written as

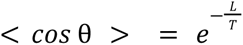

where <cos θ> is the expectation of the angle between any given initial tangent on the polymer and all tangents L distance away on the polymer where L is the contour length [34]. The persistence length is an attractive choice for characterizing mechanical properties of DNA, as it describes bending on both local and global scales, which is not fundamentally true for other metrics described below. Typical procedure for calculating static persistence length is by sequence averaging; here we use a sliding window of 50 bp. In addition to persistence length, we have chosen a variety of other geometrical metrics (**Table 1**). We describe these other metrics in the following section.

**Table 1:** Metrics used to characterize mechanical properties of DNA. Here *N* is the total length in bp of a sequence (in this case 100), *r*(*i*) is the spatial position of the *i-*th nucleotide, and *n*(*i*) is the vector normal to the *i-*th nucleotide..

| Metric Name | Equation | Units |
| --- | --- | --- |
| Static Persistence Length | $\frac{-\sum_{i=2}^N \ r(i) - r(i-1)\ }{\ln(\frac{2}{N} \sum_{i=1}^{N/2} n(i) \cdot n(i+N/2))}$ | Angstroms |
| Contour Length | $\sum_{i=2}^N \ r(i) - r(i-1)\ $ | Angstroms |
| Extension | $\ r(N) - r(1)\ $ | Angstroms |
| Global Bend | $\frac{\ r(N) \times r(\lfloor \frac{N}{2} \rfloor)\ }{\ r(N) - r(1)\ }$ | Angstroms |
| Angular Offset | $\arccos(n(1) \cdot \frac{r(N) - r(1)}{\ r(N) - r(1)\ })$ | Degrees |
| Angular Deviation | $\arccos(n(1) \cdot n(N))$ | Degrees |
| Beam Projection | $\sum_{i=2}^N \frac{r(i) - r(i-1)}{\ r(i) - r(i-1)\ } \cdot n(1)$ | Angstroms |
| Cumulative Curvature | $\sum_{i=2}^{N-1} \ r(i+1) - 2r(i) + r(i-1)\ $ | Angstroms/bp<sup>2</sup> |

*Contour length* is the length of the DNA sequence in its fully extended state and is the sum of Euclidean distances between adjacent base pairs. This metric is more descriptive when related to other metrics. *Extension* is the Euclidean distance between the first and last base pair. High extension implies a straight structure, while low values imply bending. *Global bend* is a metric implemented to account for large-scale bends in the DNA that may not be captured by focusing on vectors related only to terminal base pairs. Here, we take the distance between the base pair halfway through the sequence and the line segment that would connect the sequence’s terminal ends [35]. *Angular offset* describes the angle between the oriented normal of the first base pair and the line connecting the first and last base pair. Closely related is *angular deviation*, which describes the angle between the first base pair’s oriented normal and the last base pair’s oriented normal. *Beam projection* is a measure of how much the DNA continues in the initial direction. Stiff DNA would ideally always be extended in this direction. The contour being projected is normalized so that the contour length of the DNA is not heavily reflected in this metric. DNA with minimal beam projection would ideally reverse the initial direction as quickly as possible and then continue to be rod-like in the reverse direction. *Cumulative curvature* is a metric meant to account for local bends in DNA. Specifically, it is the sum of second differences between base pairs’ positions in space. In terms of extrema, a sequence with a zigzag structure would have a high cumulative curvature, while one with a straight structure would be low.

### Search algorithms

Even with a relatively computationally quick method of modeling, the sample space of all 100 bp DNA sequences is extremely large (4^100^ ≈ 1.6 × 10^60^distinct sequences) and practically impossible to search completely. Two approximate search algorithms were employed: a greedy algorithm [36] and a Metropolis-Hastings Algorithm with simulated annealing [37]. The greedy algorithm is a classic search algorithm and, as will be discussed in our results, ends up performing quite nicely for many parameters given the nature of the problem. The Metropolis-Hastings Algorithm with simulated annealing is another standard search algorithm that can serve as an alternative if the greedy algorithm does poorly from getting stuck in a local extrema. Since the proposal distribution can vary greatly from metric to metric, the Metropolis-Hastings algorithm was modified so that order statistics were used to evaluate where the candidate fell within the proposal distribution and determined acceptance or rejection. These algorithms were used for both the minima and the maxima of each mechanical parameter 6for both sets of DNA parametrizations. Since cgDNA+ takes a bit longer to run, we cut down on the number of sequences searched by 1/4. The overall workflow is the same for all models. The greedy algorithm was run first, followed by the Metropolis-Hastings Algorithm.

#### Greedy Algorithm

1. Initialize with a list containing [‘A’, ‘T’, ‘G’, ‘C’] and pass to algorithm.
2. For each iteration, perform the following on passed list:

a. Construct all possible 1 bp extensions (Ex. if [‘A’] was passed, create the list [‘AA’, ‘AT’, ‘AG’, ‘AC’]).
b. Calculate target metric for each entry in newly created list.
c. Sort list by target metric in ascending or descending order if search for max or min respectively.
d. Keep at most 4096 entries (1024 for cgDNA+).
e. Return list.
3. Continue iterating until target length is reached.

#### Nonparametric Metropolis Hastings Algorithm with Simulated Annealing

1. Initialize with resulting sequence from greedy algorithm, annealing factor of 33, and an acceptance percentile of 33% (more information below).
2. For each iteration:

a. Generate a candidate sequence from a proposal distribution. The proposal distribution is uniform over all sequences differing from the current sequence in the number of positions equal to the annealing factor. The annealing factor (distribution variance) is decreased every 1000 iterations from 33 to 1 at a degree of 0.5 (specifically at a schedule of 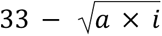 where i is iteration and a is a scaling factor).
b. Calculate the proposed sequence’s target metric of interest. Calculate the percentile of the proposed sequence’s target metric relative to the last 100 sequence target metric.
c. If the proposed sequence’s metric of interest is below the acceptance percentile reject the sequence, else accept. Ex. If the acceptance threshold is currently 50% and the proposed sequence has a parameter better than 60 of the last 100 sequences, we accept. The acceptance threshold is updated every iteration and increases from 33% to 99% exponentially (specifically the schedule is of the form *ae^i^* + *b*).
d. Repeat for 1 million iterations (250,000 for cgDNA+).

## Results and Discussion

Our results consist of the following four key takeaways. First, our results corroborate previous findings that A-tract sequences are bent [19] and poly A sequences are much straighter [12]. This not only strengthens the evidence for mechanically interesting sequences but also further validates that certain results can be derived from relatively simple models (compared to, say, molecular dynamics simulations). Second, we see evidence suggesting that extrema can be separated from each other. Specifically, we discuss how Procrustes analysis and hierarchical clustering lead to distinct clusters. Third, we sketch out a mathematical argument that explains why many of our extrema are realized by periodic sequences. Fourth, and perhaps most interestingly, the extremizing sequences have non-trivial matching sequences in current genetic databases. We detail how even our more complex discovered motifs, such as A-tracts, can obtain hits via a BLAST search. We also show that such matched wild-type sequences, although not identical to the extremizing sequences, appear to continue to display unique mechanical properties.

### Identified Extrema

The results of the extrema search reveal specific common motifs (**Table 2**). A more detailed table can be found in the SI along with metric values (**SI Table 1**). Our results agree well with previous results in validating the stiffness of poly A sequences and the bent structure of A tracts which both appear in our extrema. **Figure 1** displays what such sequences look like.

**Figure 1:**
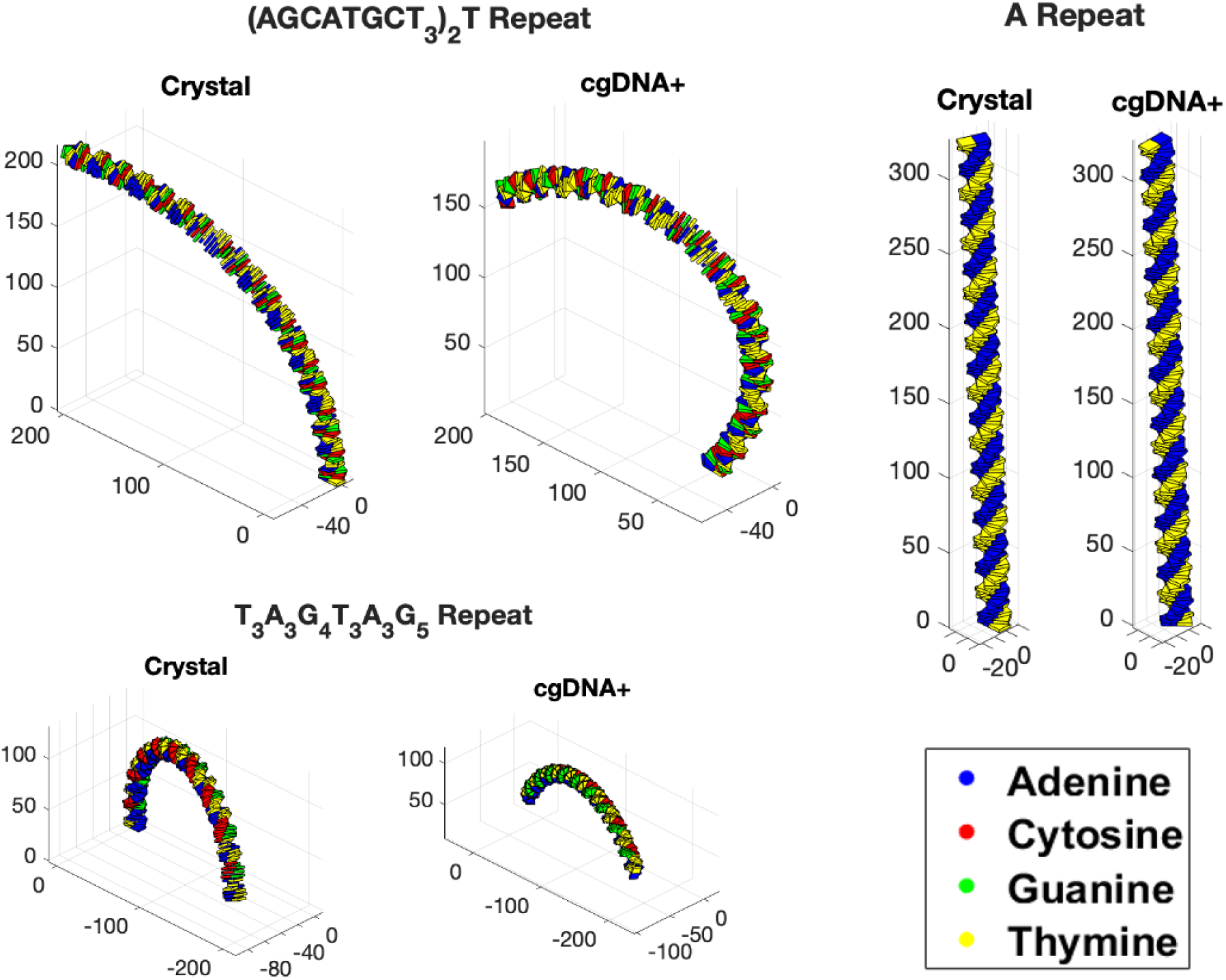
Selected 100 bp long DNA sequences composed of motifs common in found extrema. Coordinate space is in Angstroms. Both the sequences parametrized by X-ray crystallography derived parameters and cgDNA+ are displayed. The crystal sequence begins from the origin while the cgDNA+ sequence is aligned in space to the crystal sequence via Procrustes analysis. The colors of the base pairs are blue for adenine, yellow for thymine, green for guanine, and red for cytosine. One can see that (A) repeat is quite straight in both crystal and cgDNA+. The bent sequences are similar in both the crystal parametrization and the cgDNA+ model with each sequence being more bent with its own extremizing sequence.

**Table 2:** Selected motifs from extrema search. Only motifs that had extrema in multiple metrics are listed. Other motifs can be seen in **SI Table 1**. The motif is on the left hand side and the metrics the sequence is an extrema in are on the right hand side. These motifs are representative of found extrema sequences. Actual extrema may differ from the repeats though be of a similar form.

| Motifs of Interest | Extrema Parameters |
| --- | --- |
| (A) Repeat | Greatest Static Persistence Length (All)<br>Greatest Beam Projection (Crystal/Molecular)<br>Least Cumulative Curvature (All) |
| (T<sub>3</sub>A<sub>3</sub>G<sub>4</sub>T<sub>3</sub>A<sub>3</sub>G<sub>5</sub>) Repeat | Least Static Persistence Length (Crystal)<br>Least Extension (Crystal)<br>Greatest Global Bend (Crystal)<br>Greatest Angular Offset (Crystal)<br>Greatest Angular Deviation (Crystal)<br>Least Beam Projection (Crystal) |
| (AGCATGCT<sub>3</sub>)<sub>2</sub>T Repeat | Least Static Persistence Length (cgDNA+)<br>Least Extension (cgDNA+)<br>Greatest Angular Offset (cgDNA+)<br>Greatest Angular Deviation (cgDNA+)<br>Least Beam Projection (cgDNA+) |
| (CATG<sub>3</sub>A<sub>5</sub>) Repeat | Least Static Persistence Length (Molecular)<br>Least Extension (Molecular)<br>Greatest Global Bend (Molecular)<br>Greatest Angular Offset (Molecular)<br>Greatest Angular Deviation (Molecular)<br>Least Beam Projection (Molecular) |

When exploring extrema for metrics where such values would imply straightness, we see that poly A sequences appear frequently in all models. For greatest static persistence length, the actual sequences found consisted of a very T heavy sequence in the first half with a very A heavy sequence in the second half (**SI Table 1**). It is likely that the deviations from exact poly A for the actual extrema arise from either the imperfect nature of the static persistence length metric used where tangent correlations are calculated consistently at 50 base pairs apart, as an artifact of finite length of the extremized sequence, extremizing ambiguity from the fact mechanical and structural properties of two complementary sequences are identical, or from a minor benefit obtained at the polyT-polyA interface. Regardless, the actual poly A sequence still has an exceptionally high static persistence length (**SI Table 1**).

We used multiple metrics to avoid the shortcomings of single metrics and observed poly A sequences appear for more metrics such as high beam projection and low cumulative curvature. A very high beam projection implies that repetition of A causes the DNA sequence to maintain its initial direction very tightly. Low cumulative curvature suggests little change from tangent to tangent in poly A sequences. This indicates that our results with poly A sequences are consistent with other works examining persistence lengths of DNA [12].

Some other metrics that may be associated with straightness, such as least angular deviation, least angular offset, and low global bend, were not very informative. Since these metrics depend on at most three points, the majority of the sequence can be used to align points in just a couple of base pairs in a straight line, which minimizes these metrics. While these metrics are more useful for analyzing more curved and bent sequences, the results served as a null model when addressing prevalence in biological data, as no evidence of these sequences occurring naturally was found (**SI Table 1**).

Much of the extrema for metrics characterizing deformed structures are realized by A-tract-like structures. When searching the sequence space parameterized by X-ray crystallography data, the motif that consistently appears in various metrics implying bending and curving is T_3_A_3_G_4_T_3_A_3_G_5_ repeated (**Table 2, SI Table 1**) . This sequence is similar to an A-tract, but A-tracts are typically defined as having four or more A’s without a TA step [21], while here we see three T’s and three A’s. The entire motif spans 21 bps, which is in phase with the standard DNA helicity of 10-10.5 bp [38]. Although the actual found extrema are not precisely this repeated motif, repeating this motif recreates near-extreme values (**SI Table 1**). Bent extrema for cgDNA+ tended toward the motif of (AGCATGCT_3_)_2_T (or in its reverse complementary form A(A_3_GCATGCT)_2_. Like extrema for the X-ray data, the motif contains A runs and is a 21 bp motif. This motif shows up slightly more weakly than the crystal motif, potentially a byproduct of a more complex search space given cgDNA+ takes into account nonlocal effects and the reduced search algorithm. However, repeating the motif provides a quite bent structure and interesting BLAST results, as will be discussed. As for molecular dynamics-derived values, we again find A-tract-like sequences as extrema. These A-tracts are more diverse than those in the extrema of the crystal parameters, but they can be roughly characterized as the motif CATG_3_A_5_ which is interestingly a 10 bp repeat rather than a 21 bp repeat (**SI Table 1**).

The remaining extrema sequences also display extrema characteristics, primarily leaning towards stiffness through the use of short repetitive and periodic motifs. Repetition and periodicity have proven to be emblematic of almost all extrema we have found. In later sections, we present a mathematical argument as to why this is unlikely to be coincidental.

### Structure of Sequence Space

Before discussing why extrema are periodic or how they can appear via BLAST searches, it is worth examining how distinct extrema actually are. Specifically, to what extent do the extrema differ geometrically from the rest of the sample space? To compare the DNA structures produced by the sequences, we employed Procrustes analysis [39] to assess sequence shapes. Procrustes analysis optimally aligns the 3D structures and then calculates the standardized sum of squared distance between corresponding base pair centers as a metric for shape similarity. To visualize the resulting distances, hierarchical clustering with complete linkage was performed.

The following sequences were used to construct the resulting dendrogram (**Figure 2**). First, we used each motif mentioned in **Table 2**, repeated to 100 bps. Both the version parametrized by crystallography-derived parameters and the version using cgDNA+ were used. This resulted in a total of six structures identical to the six shown in **Figure 1**. Next, we chose to utilize random sequences with AT or GC bias. AT or GC bias means that AT (or GC) is four times as likely to appear in a random sequence than GC (or AT). 50 sequences were randomly generated for each bias and for each model. Finally, all of these sequences needed to be compared to random sequences with no GC/AT bias. For these sequences, when comparisons were made between a random and another sequence, the Procrustes distance was calculated 50 times with the random sequence being randomly generated each time, and the average was used in construction of the dendrogram.

**Figure 2:**
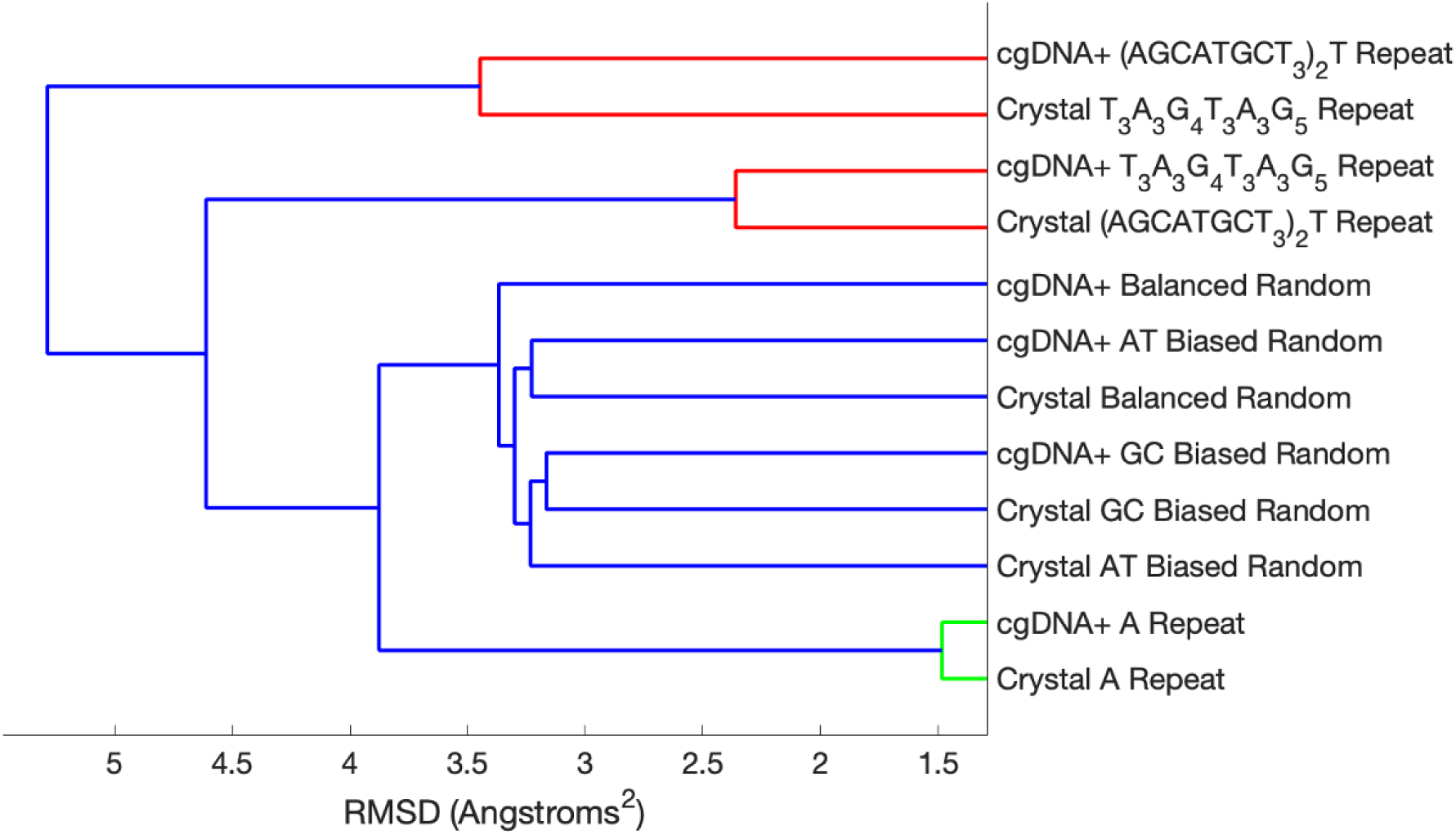
Hierarchical clustering of various DNA sequences as compared by Procrustes distance separates out extrema. The labels can be read in the following manner. The first word is either cgDNA+ or crystal depending on which set of parameters was used to generate the DNA structure via the Cambridge Scheme. We included all motifs mentioned in **Table 2** repeated to 100 bps in length. Random sequences are taken to be at the average of 50 randomly generated sequences Procrustes distance to compared sequences. Units are in Angstroms squared.

We observe that the A tracts and poly A sequences separate out well. We see that extremizing motifs parametrized with their respective models are more similar to each other than the same sequence between two models. The intermediate grouping contains the tested non-extrema sequences which all have very similar distances between themselves and no clear smaller groupings. We have also recorded nonparametric summary statistics for each metric over 10,000 randomly generated 100 bp sequences (**Table S1**). These numbers further demonstrate that the found extrema significantly deviate from random sequences.

### Periodic Nature of Extrema

For certain types of metrics we present a mathematical argument suggesting that the extrema for that metric will be periodic DNA sequences. A non-technical argument is provided here, while a rigorous proof can be found in the supplementary information (**SI Text**). We use the method of considering DNA sequences on a directed graph. Such an approach has been used before to identify how genomic sequences could be modified to best accommodate nucleosomes via a shortest path algorithm while keeping genetic information intact [40]. Here we use a graph theory approach to characterize what type of sequences mechanical extrema must be.

Consider DNA as a chain made of a finite number of distinct segments, where each segment contributes to a cumulative metric linearly. The simplest example is the contour length, which is a sum of the contour lengths of individual base pair steps. Since not all base steps are independent of each other (e.g. AT step cannot be followed by GC step), it is not possible to repeat the step with the largest contour length in order to obtain an extremum.

Now let’s examine the challenge of determining an infinite DNA sequence with the maximum possible contour length. This problem can be conceptualized as a procedure to discover the heaviest path on a directed graph (**Figure 3**). For any given vertex in the graph, there is a strategy to maximize the contour length: follow the path along the edge with the highest weight. However, an important aspect to remember is that the number of edges is finite. This means that, eventually, we will circle back to a vertex previously visited. Even though we may find ourselves back at the starting point, the graph structure remains unchanged. Consequently, the optimal strategy also remains constant, and we will follow the same path again, leading us once more to the current vertex. This cyclical process implies that the sequence with the greatest possible contour length must inherently be periodic. This can be generalized to any number of finite vertices where each vertex can represent a finite number of consecutive base pairs.

**Figure 3:**
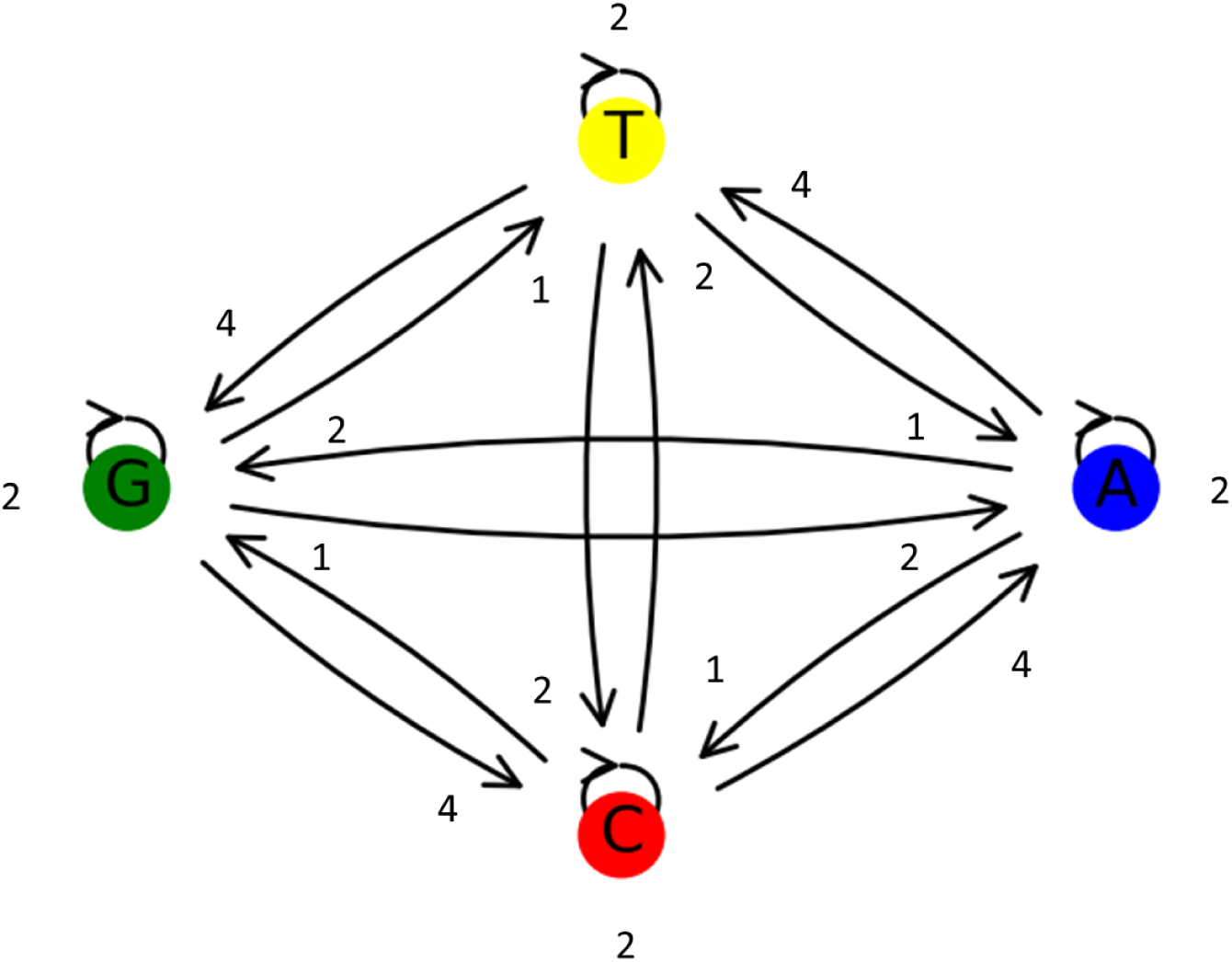
DNA sequences can be visualized as a walk on a directed graph. An example of how to maximize a metric based on DNA sequence can be visualized as a connected directed graph. Each vertex represents a nucleotide while each edge represents a nucleotide step. The edges carry weights given here as arbitrary integers.

In practice, there are some limitations to consider. Firstly, DNA sequences are finite, which means it cannot be conclusively demonstrated that a finite sequence has to be periodic. This is because the final segment might be unique, as there is no mandatory subsequent segment. A similar situation could occur for an initial segment. This accounts for why some of the observed extreme cases are just a few base pairs short of being truly periodic. However, finite length should not have a significant practical effect, as long sequences will primarily consist of periodic structures, with potential exceptions at the beginning and the end.

More pertinent is the fact that segments may not independently contribute to the desired metric. However, given that the persistence length for DNA is not infinite, distinct segments should arise eventually for very long sequences. There is also the natural helicity of DNA, which could allow for initial conditions to be reset to an extent (as in the sequence in one or two turns of the helix is independent of the sequence in other turns of the helix). This could explain how motifs of size 11 and 22 were found, which are in phase with the natural helicity of DNA.

### Biological Prevalence of Periodic Sequences and Structural Extrema

To assess the frequency of occurrence of the identified sequences in living organisms, we conducted BLAST searches [41] with some modifications to the default settings of the nucleotide BLASTn program. We disabled low complexity filtering as we are interested in low complexity regions. The searches were performed against the standard database, which includes RNA transcripts that may have undergone post-transcriptional modifications in eukaryotes. We opted to include these transcripts, as they are prevalent in most databases, primarily derive from the original DNA, and could potentially exhibit similar mechanical behavior to RNA sequences. For our motifs, we extended them up to 500 bp when performing BLAST searches. All searches were carried out in November 2023, and it should be noted that the BLAST database is continually expanding, potentially leading to new matches in the future. We used the gap penalty of 5 for existence and 2 for extension and mismatch penalties 2, -3. To evaluate the quality of the BLAST search results, we considered two metrics: the *number of hits* and the *bit score of the top hit*. The hits are based on an expected statistical significance threshold of 5%. We chose to only count hits that had a p-value below 10^-50^ (expect threshold=10^-50^ ). The bit score, measured on a log-2 scale, represents the expected size of a database in which a hit would occur by random chance. For instance, a bit score of 500 corresponds to an expected database size of 2^500^.

### Periodic Sequences

By repeating randomly generated DNA sequences of 11 or 21 bp up to 500 bp, we obtained multiple high-scoring BLAST hits. This finding is not entirely surprising, considering the prevalence of repetitive DNA [42] and the possible increased ease of alignment for sequences with existing repetitive motifs. However, the sheer number and high quality of hits are rather unexpected, especially given the length of the repeats. For 20 randomly generated motifs of 11 bp extended to 500 bp, the average number of hits was 57.8 (variance: 5,268), and the average top bit score was 545.25 (variance: 15,445). The Spearman Rank Correlation between these parameters was 0.576. For 20 randomly generated motifs of 21 bp extended to 500 bp, the average number of hits was 10.9 (variance: 173.7), and the average top bit score was 308.2 (variance: 8,916). The Spearman Rank Correlation between these parameters was 0.716.

The high variance observed is notable; if motif composition were inconsequential, the data might follow a Poisson distribution, particularly for the number of hits, implying equal mean and variance. However, this is not the case. The cause of the high variance is unclear and suggests not all motifs are equally biologically present. Perhaps the most straightforward possibility would be that more complicated motifs are less prevalent. To briefly examine this, we can calculate the Spearman Rank Coefficient of the number of hits and the top bit score to the Shannon entropy of the sequence (as defined by nucleotide frequency). Higher Shannon entropy could be interpreted as a potentially more complex sequence. For 11 bp, the Spearman Rank

Coefficient is -0.193 for number of hits and 0.205 for top hit score. For 21 bp, the Spearman Rank Coefficient is -0.258 for number of hits and -0.262 for top hit score. This suggests sequence simplicity could make motifs more likely to be repeated but is likely not on its own a sufficient explanation, necessitating future research. Individual sample data is available in the SI (**SI Table 2**, **SI Table 3**).

### Extremizing Sequences

The next step is to evaluate the performance of our proposed motifs, specifically those in **Table 2**. Including transcripts in the search for a 500 bp repeat of adenine does impact the results, as eukaryotic post-transcriptional modifications often involve the addition of poly A tails [43] to protect mRNA from degradation. This may not be a coincidence. However, since our focus is on DNA, we explored DNA sequences by performing a BLAST search for a 500 bp A tract on the human reference genome. We restricted ourselves to the human genome as attempting to BLAST the standard database through an error (RAM usage exceeded). We found 184 hits, with the highest-scoring hit at 579 bits, located on chromosome 2. The subsequent two best hits scored 255 bits and 246 bits, respectively. Given that high-scoring hits are present within the human genome, it is highly probable that such long A tracts also occur in other organisms’ genomes.

For the more bent extrema, we observed the following results. For the (T_3_A_3_G_4_T_3_A_3_G_5_) repeat there were 141 hits with a top hit score of 519. The top hit was found in the genome of *Branchellion lobata* (Species of Leech). The next two best hit scores were 511 bits found in the genome of *Blennius ocellaris* (Butterfly Blenny) and 488 bits in the genome of *Sphaeramia orbicularis* (Orbiculate Cardinalfish). When compared to the 20 randomly generated motifs of size 21 bp, this sequence has a greater number of hits than 20/20 and has a higher top hit score than 19/20.

For the (AGCATGCT_3_)_2_T motif there were 438 hits with a top hit score of 411 bits. The top hit was found in the genome of *Gobio gobio* (Gudgeon). The next two best hit scores were 391 bits found in the genome of *Gobius niger* (Black Goby) and 386 bits in the genome of *Danio aesculapii* (Species of Carp). When compared to the 20 randomly generated motifs of size 21 bp, this sequence has a greater number of hits than 20/20 of those sequences and has a higher top bit score than 18/20.

For the (CATG_3_A_5_) repeat there were 102 hits with a top hit score of 515 bits. The top hit was found in the genome of *Erithacus rubecula* (European Robin). The next two best hit scores were 464 bits found in the genome of *Taurulus bubalis* (Species of Sculpin) and 460 bits in the genome of *Psylliodes chrysocephala* (Species of Flea Beetle). When compared to the 20 randomly generated motifs of size 11 bp, this sequence has a greater number of hits than 16/20 of those sequences and has a higher top bit score than 12/20.

We also chose to compare our results to the motif used for A tracts in reference [10] which were in turn motifs used in experimental work [44]. This motif is of the form (A_6_CG_2_CA_6_CG_3_C). There were 51 hits with a top hit score of 593 bits. The top hit was found in the genome of *Psittacula echo* (Mauritius parakeet). The next two best species unique (there were a couple of high quality hits in different chromosomes of *Zonotrichia albicollis)* hit scores were 582 bits found in the genome of *Zonotrichia albicollis* (Species of Snakefly) and 369 bits in the genome of *Gobius niger* (Black Goby). When compared to the 20 randomly generated motifs of size 21 bp, this sequence has a greater number of hits than 19/20 of those sequences and has a higher top bit score than 20/20.

We thus find that not all A tract sequences are necessarily equivalent. The best scoring hit came from the experimentally used A-tract motif of (A_6_CG_2_CA_6_CG_3_C) though this motif also had the smallest number of total hits. On the other hand the cgDNA+ motif (AGCATGCT_3_)_2_T had the lowest scoring top hit but by far the greatest number of hits. The majority of these hits came from chromosomes of various strains of *Danio rerio* (Zebrafish) and related species. This suggests that not only is a predicted extrema motif present in real organisms but potentially conserved. This could be the case for some of the other motifs but they may appear in species that currently have less diversity in their available genomes as they are not model organisms like zebrafish. It is likely that the motifs we have tested are not completely optimal due to limitations in searching the sample space along with the imperfection of models and metrics. Furthermore, evolution itself is not necessarily optimal and good enough A-tracts may often end up being chosen. However, we do observe enrichment for our known mechanically interesting sequences in terms of the number of hits and top hit scores, particularly for the 21 bp motifs.

Another question is whether these top hitting sequences actually look bent and curved. Indeed we see that these biological sequences best matching the repeated motifs display curving and bending properties (**Figure 4**). The (T_3_A_3_G_4_T_3_A_3_G_5_) repeat hit displays something of a secondary helical structure with both forms of parameterizations, though more so with the crystal structure. The (A_6_CG_2_CA_6_CG_3_C) repeat hit creates a loop with both the crystal model and the cgDNA+ model. Finally, (AGCATGCT_3_)_2_T motif displays a couple of major overarching bends. The random reference sequence shows that the average random 500 bp sequence will be somewhat straight with the occasional kink.

**Figure 4:**
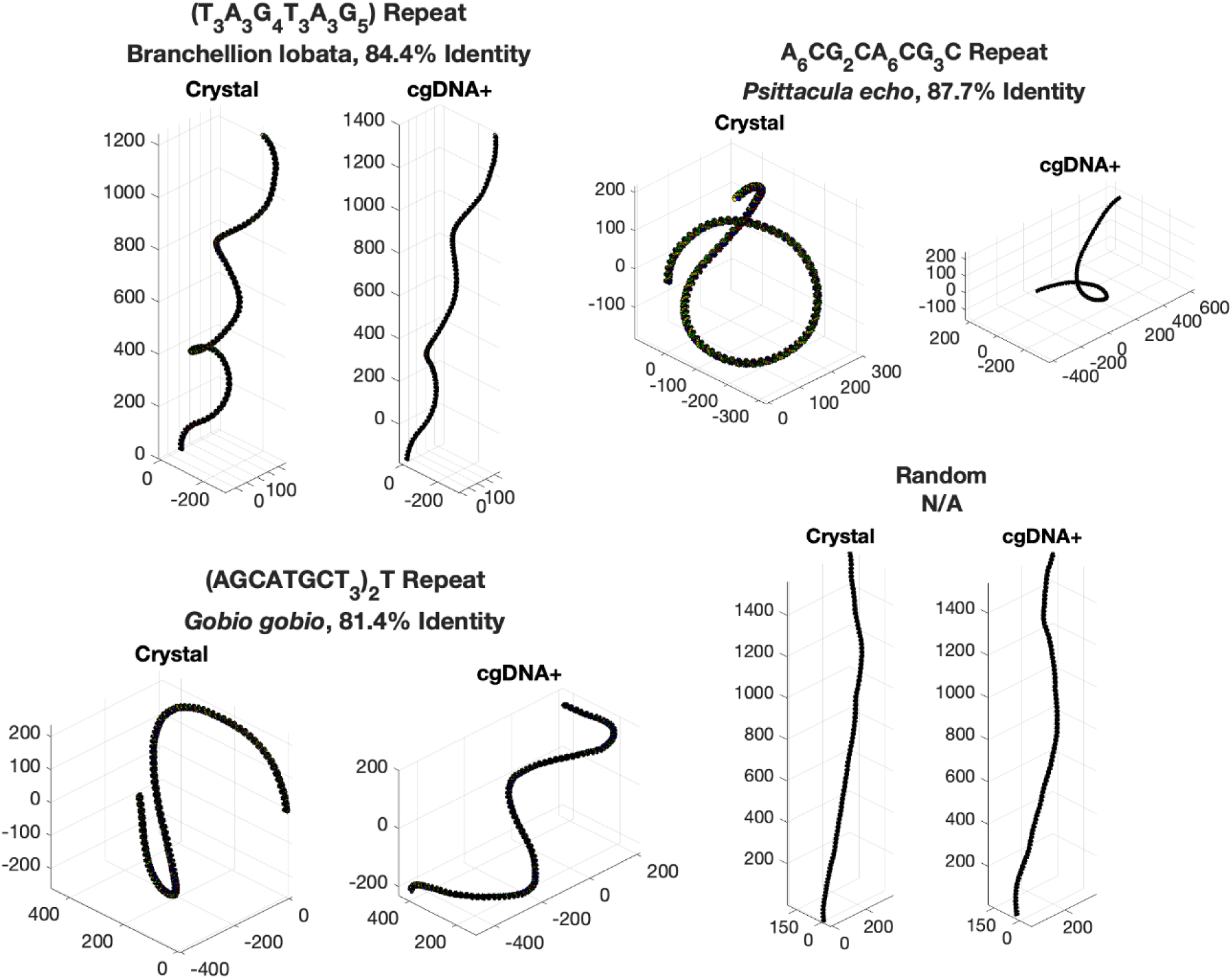
Sequences found during BLAST searches for repeats display loops and bends. Coordinate space is in Angstroms. The specific sequences can be found in the supplementary information **Table S4**. All of the sequences are found biologically with the exception of the random sequence which is randomly generated DNA of 500 bp length for reference. Both the sequence parametrized by crystal values and molecular values are shown. Below the name of the repeated motif, the species the sequence originated in along with the percent identity of the aligned sequence to the original motif is given.

We also conducted a brief analysis of all 100 bp sequences as obtained directly from the search algorithm rather than after motif simplification and extension to 500 bps. Many of the sequences returned hits when MegaBLAST was employed, using default databases and parameters except for unmasking low complexity regions. The crystal and molecular datasets obtained more MegaBLAST hits, though as mentioned earlier this can stem from the search algorithms not being as effective in the more complex search space coupled with the slightly reduced algorithm. Other notable exceptions where no hits were found included sequences with the least angular offset, least angular deviation, and least global bend—the three metrics mentioned earlier as ineffective for searching due to their optimization relying on at most three-point alignments. We provide the top hit scores and number of hits for these sequences (**SI Table 1**).

## Conclusions

The original goal of this work was to characterize what sequences make up the mechanical extrema of DNA sequence space. In the process of doing this, we discovered that almost all of the extrema displayed a degree of periodicity. These extrema were distinct from random DNA sequences to a point where it appears feasible they could have unique properties. We also were able to analytically prove that under certain assumptions, repetitive DNA sequences would have to be the extrema for a variety of mechanical properties. Finally, we were able to show these repeats matched well against existing biological sequences and even though the wild type sequences were not identical to the extremal sequences, they still displayed high levels of bending in the case of A tracts or stiffness in the case of poly A.

We propose that the prevalence of repetitive DNA [25] may be partly attributed to the unique mechanical properties that these sequences can possess, which would not be present in random or seemingly random DNA sequences. Properties that are the sum of their parts inherently have extrema that result from maximizing a part and then repeating it. As we showed, this should hold even in situations where choosing one part constrains the choice of the next part, as long as one can separate parts into unique segments.

The functional implications of these repetitive sequences with unique properties remain an open question. As demonstrated in **Figure 4**, some sequences appear highly compact, suggesting a potential role in DNA compaction. The curving may also facilitate interactions with proteins, while stiff DNA sequences could have the opposite effect. Understanding these properties is important as the genome is well known to be regulated via looping and compaction [2]. Another possibility is that these DNA sequences are more easily propagated due to effects on DNA replication. Furthermore, the transcripts may also possess distinct properties, as they too would consist of repeated elements.

Another aspect to consider is the level of conservation. The (AGCATGCT_3_)_2_T motif hints at the possibility of conservation in zebrafish though it is in no way conclusive. Beyond that motif, there seemed to be limited sharing of these sequences between similar organisms, except for their apparent eukaryotic specificity. However, it is worth noting that these repeats might be more robust to certain changes compared to proteins, rendering standard methods for detecting conservation insufficient. The results we obtained were not exact repeats, and it is possible that as long as a sequence remains highly curved, most changes in sequence result in near-neutral selection levels. Some motifs may be more prevalent because they are closer to the true extrema, but selection pressure could be weak enough upon approach to the extrema that various sequences could function effectively.

The models of DNA energetics used in our analysis are based on a rigid base pair model. Such a model is approximate at the benefit of speed being able to execute on a desktop computer in well under a second. As computing resources become more available and better algorithms are developed, coarse-grained models [45] might become more computationally feasible. However, as of now coarse grained models are not at level of less than a second computation on standard computers.

That being said, the rigid base pair model alone with different parameter sets provided extrema expected from experimental results and sequences similar to these extrema were able to be found in genomes. While more advanced models could potentially find even better matching sequences or perform better with longer motifs, it still stands that repeated motifs of not insignificant length (11-21 bp) can be found throughout genomes and that, at least theoretically, mechanical extrema will be periodic sequences.

Thus, we suggest future work in this area should focus on a more comprehensive characterization of these repeats throughout known biological databases. Given these repeats exist, it could be very interesting to instead search through repeats in the genetic database and analyze their mechanical properties. A more in-depth analysis should be conducted to determine how the appearance of repeats may deviate from expectations, and theoretical frameworks should be developed to calculate expected distributions, as opposed to the initial assumption of a Poisson distribution. Furthermore, it would be valuable to identify the genomic locations where specific types of repeats occur (e.g., near genes, centromeres, telomeres, etc.). While it is already known that repeats occur in some of these locations, such as telomeres [46], it would be informative to identify the most frequently used repeats and investigate their unique structures.

Another extension would be considering the effect of methylation which can now be done with newer models like cgDNA+ [24]. However, we chose to not include methylation in our search results for a few reasons. First and foremost, the NCBI BLAST database does not support methylation so we cannot actually easily search for methylated sequences in biological databases. Second, DNA methylation rates vary among species. All domains of life use the four standard nucleobases while DNA methylation is near nonexistent in some organisms [47].

Finally, DNA methylation is an epigenetic modification and changes more quickly than the underlying DNA sequence itself adding additional complications to ascribing properties to a given sequence. Thus, we found methylation to be outside the scope of our current project. However, DNA methylation certainly plays a role in the mechanics of DNA and hopefully future research can develop a framework to include it.

Overall, we hope this work will help stimulate research into both further addressing DNA repeats the so-called ‘dark matter’ of DNA while simultaneously moving towards a better understanding of the ‘mechanical code’ of DNA.

### Code Availability

Code and parameters can be found at https://github.com/PaulTorrillo/DNAMechanicalExtrema/tree/main

## Supplementary Information

**Table S1:**
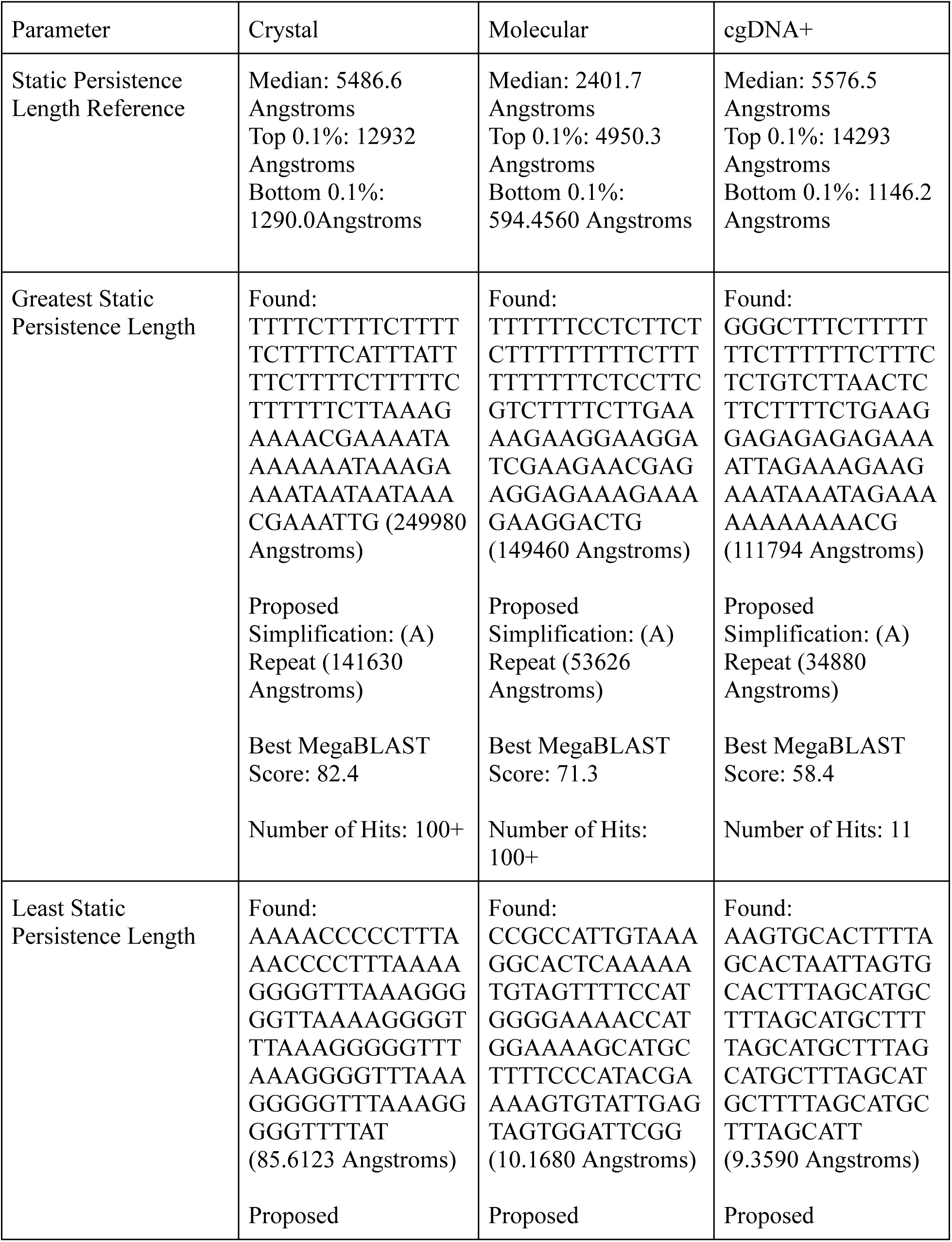
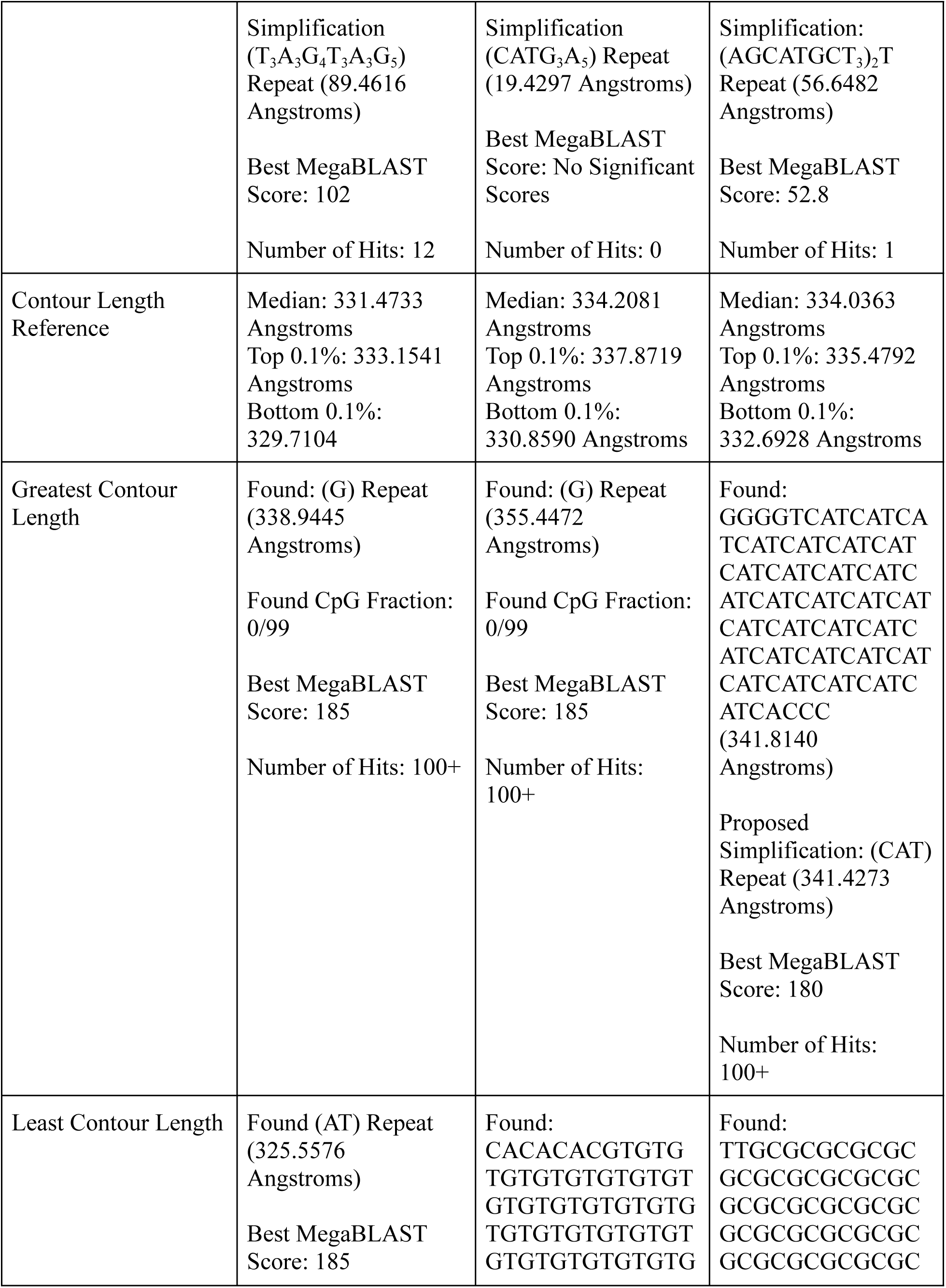
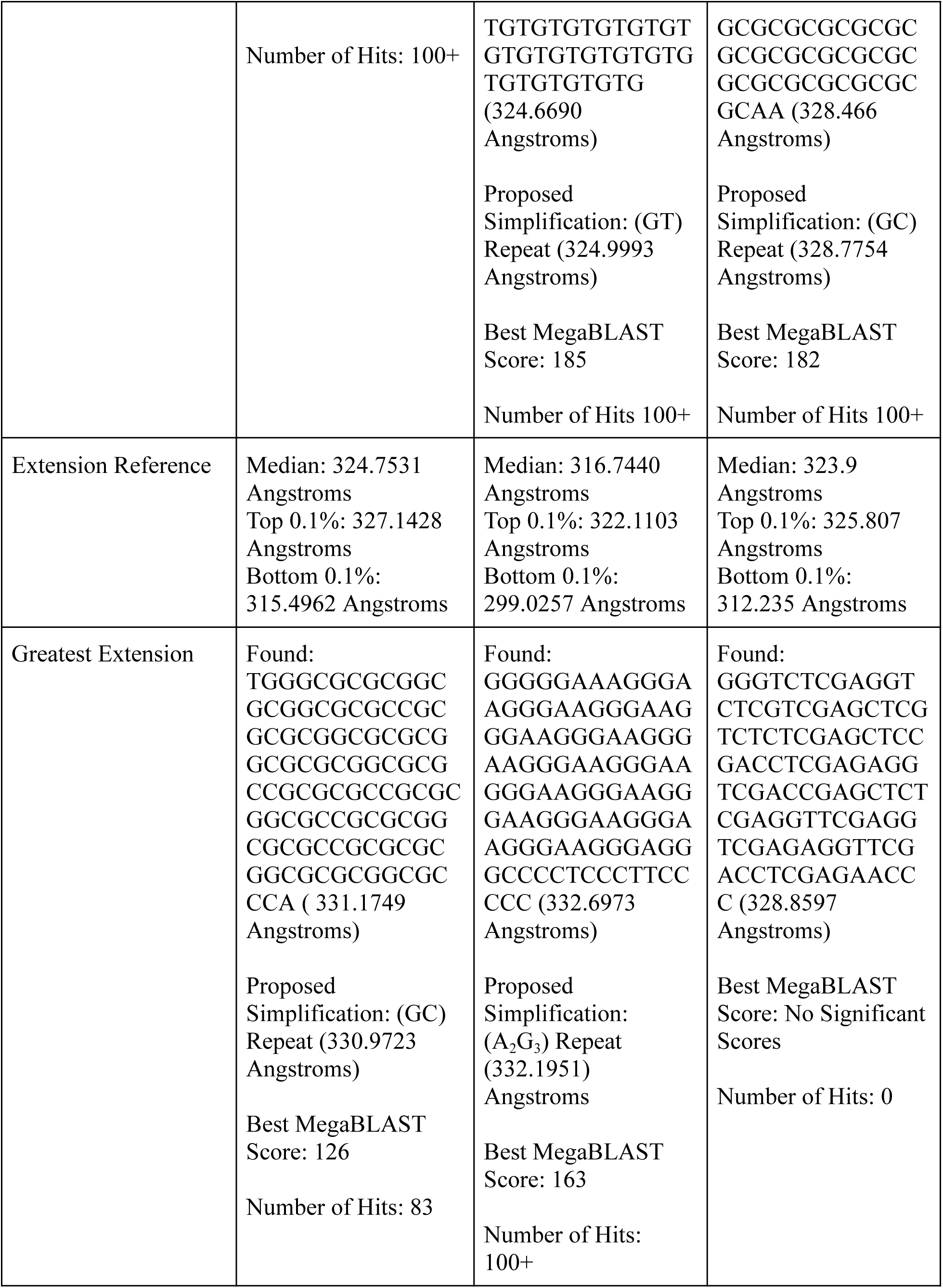
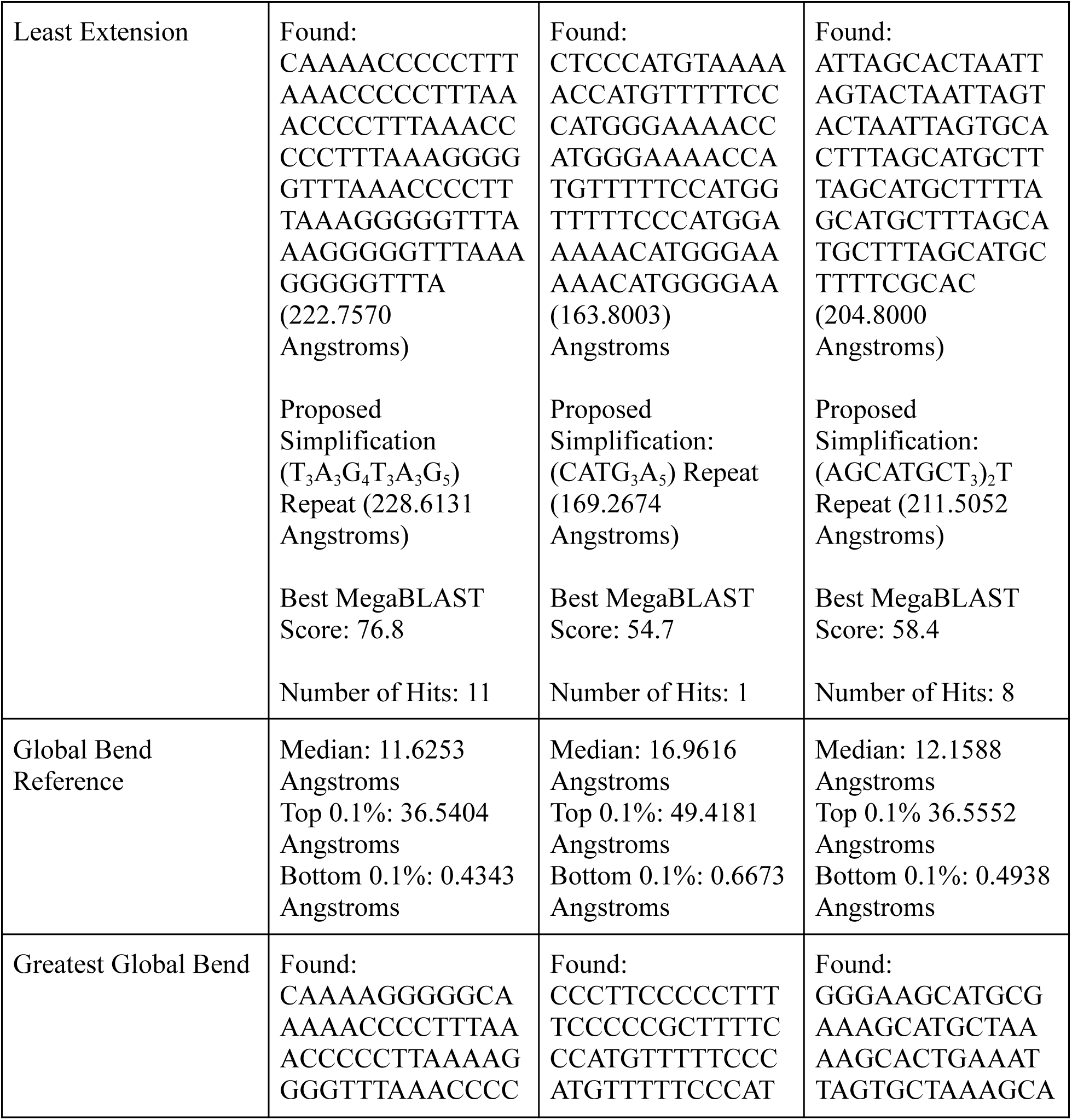
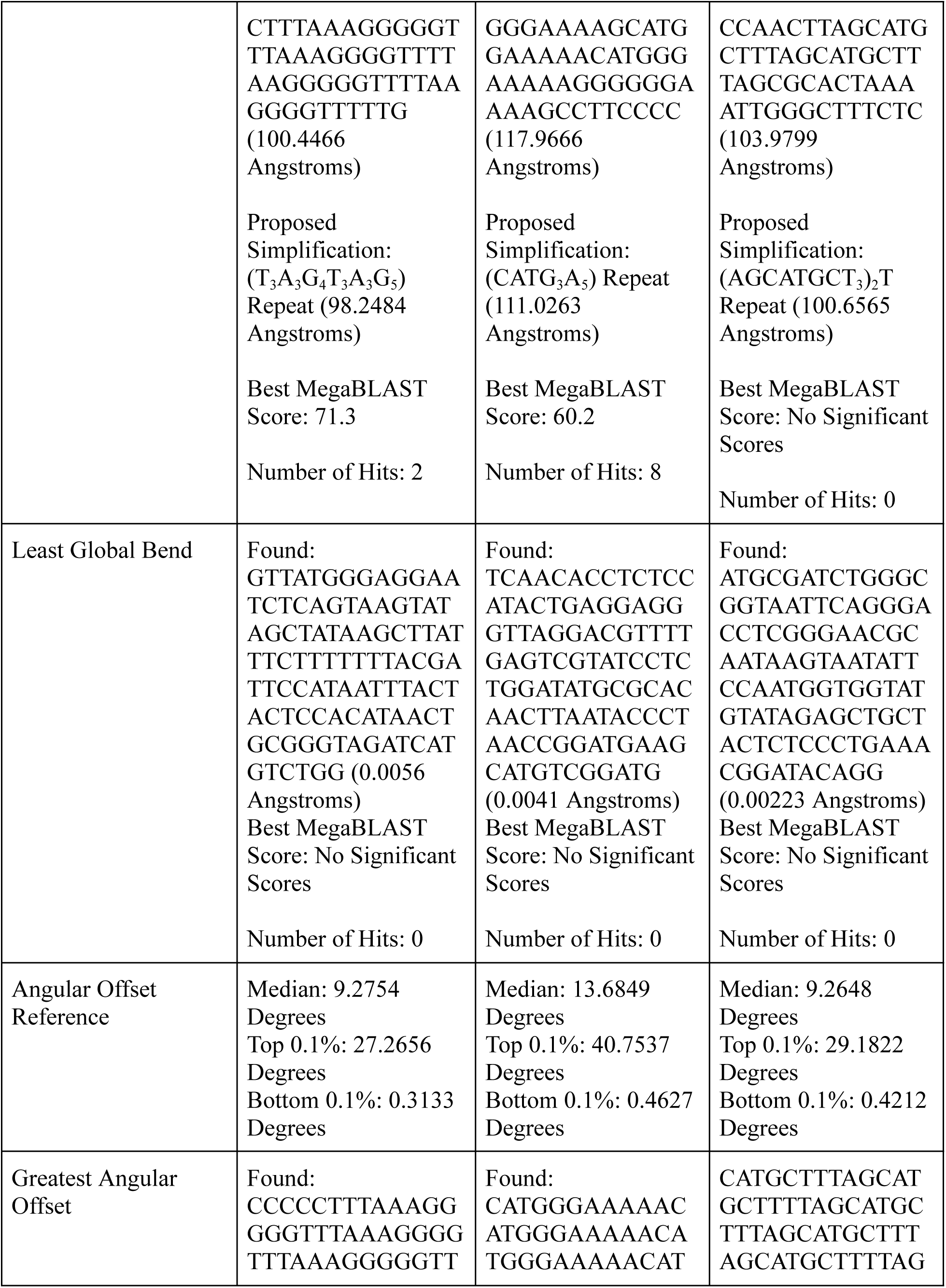
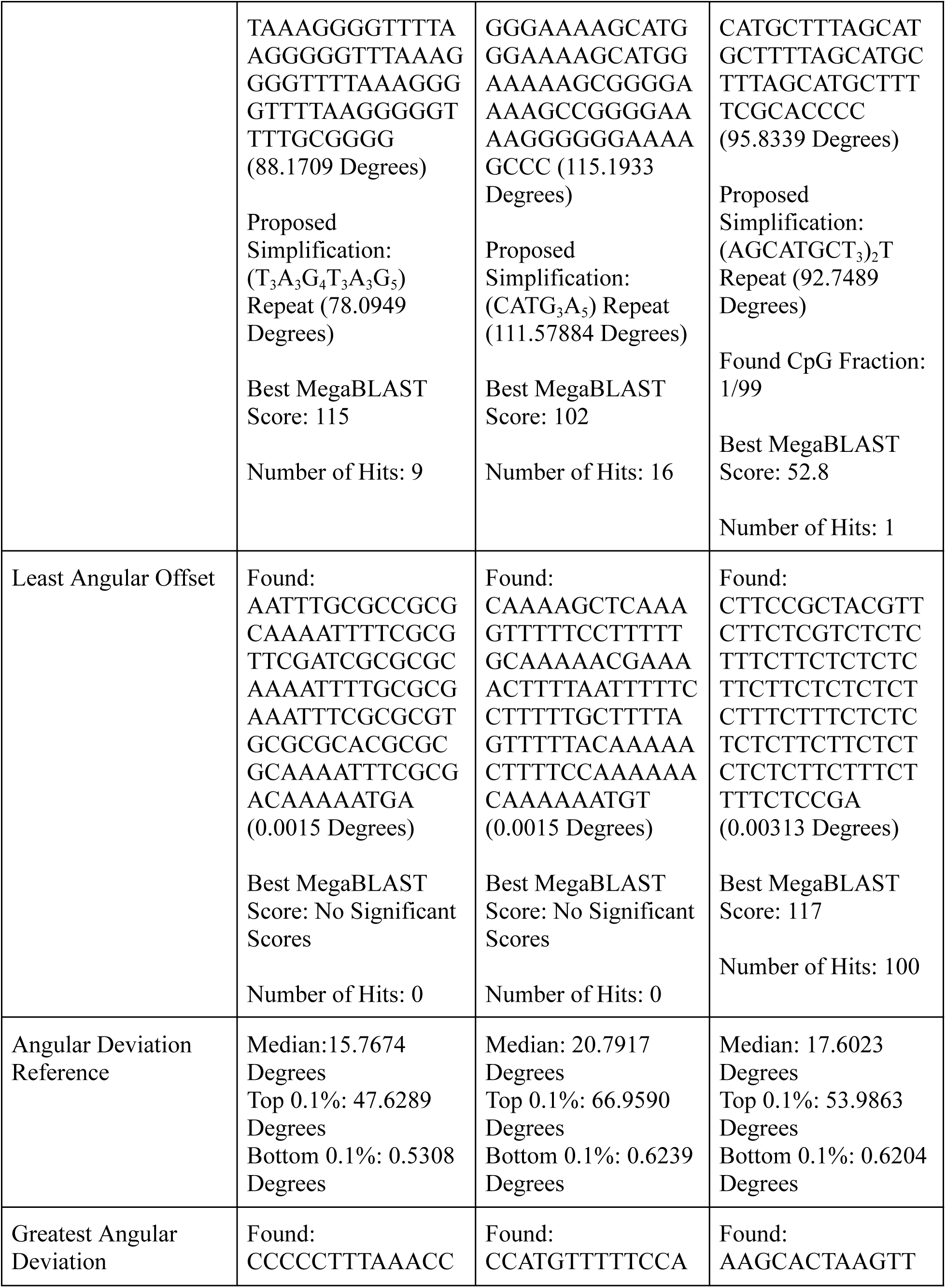
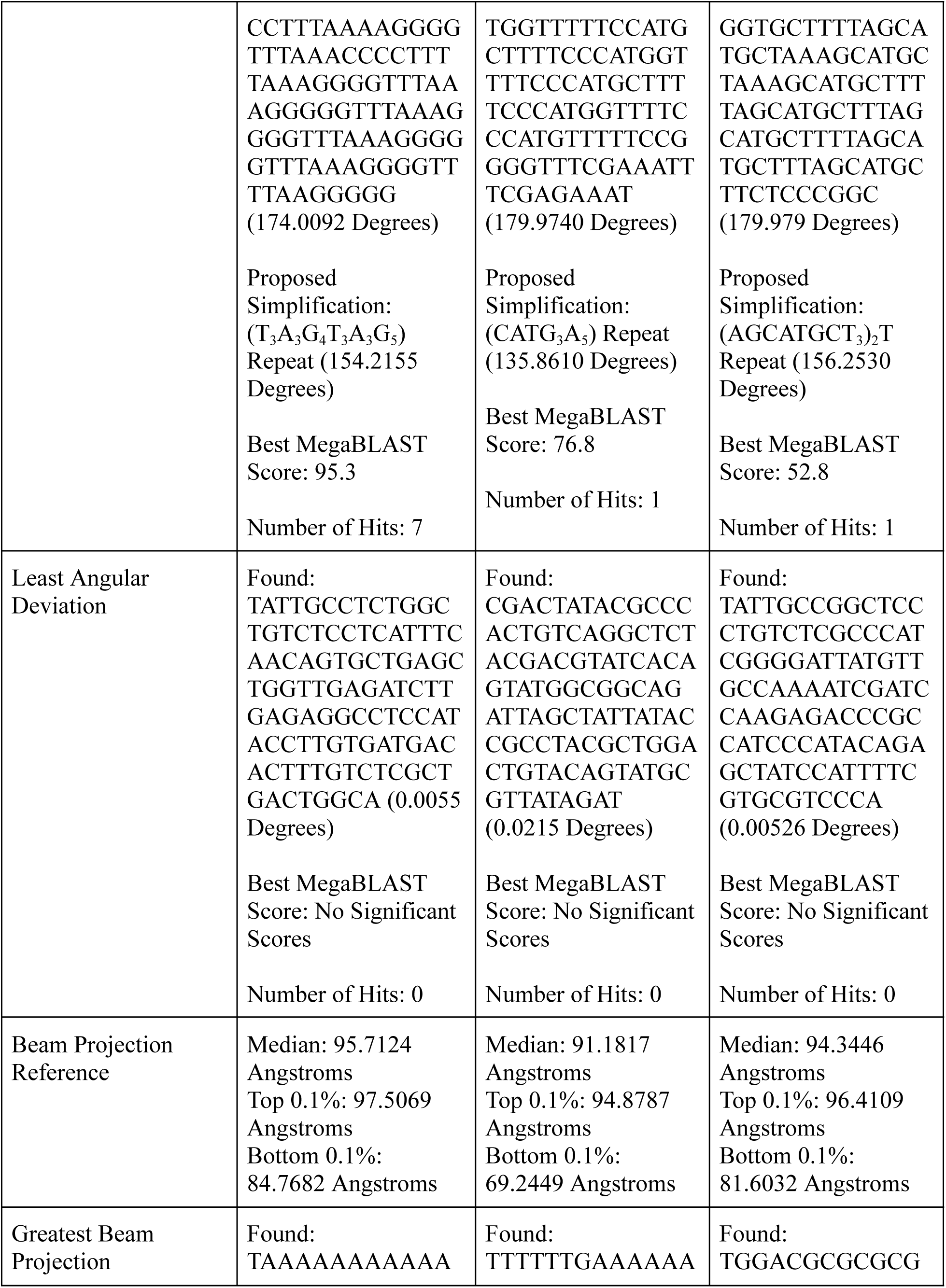
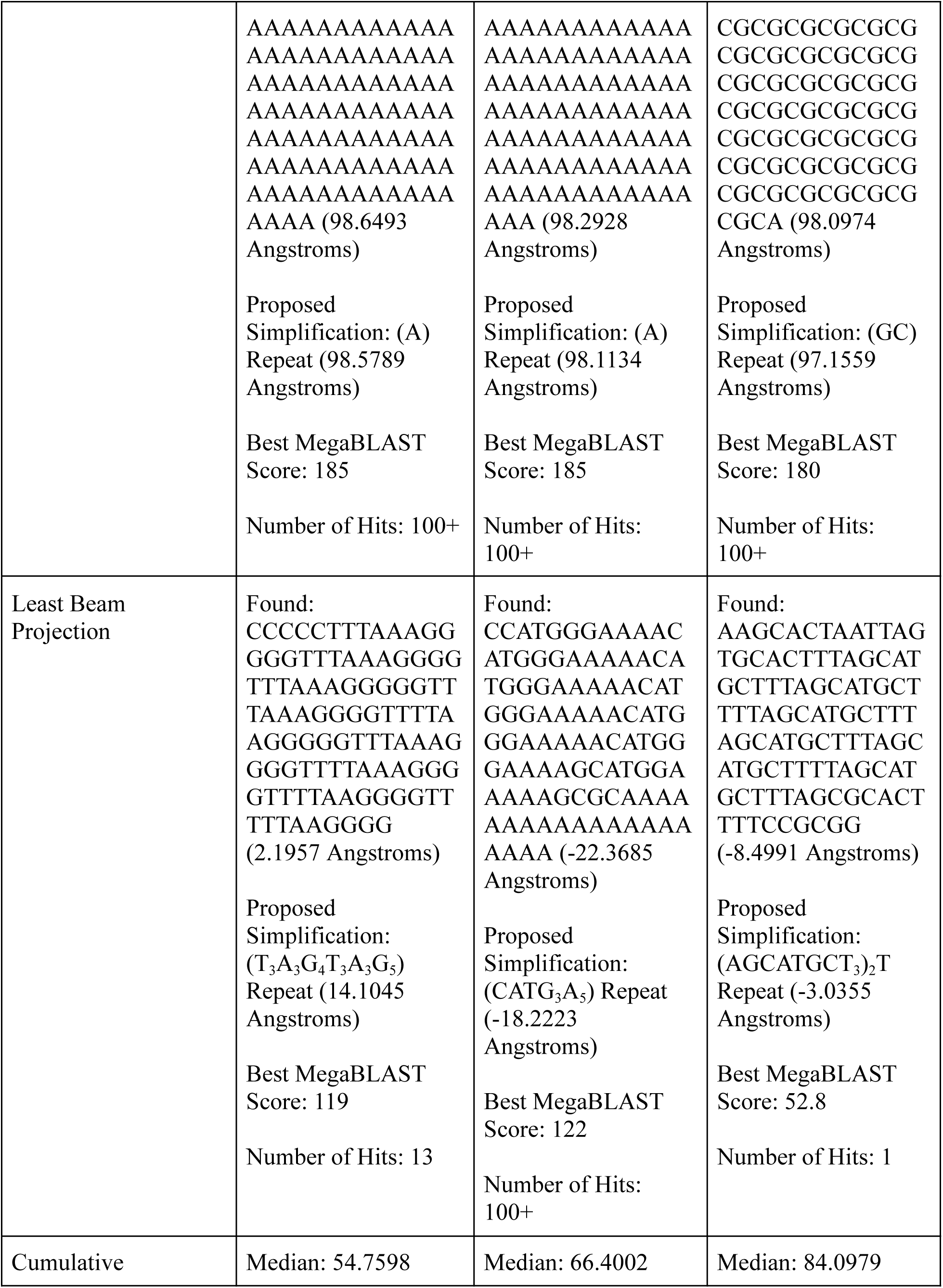
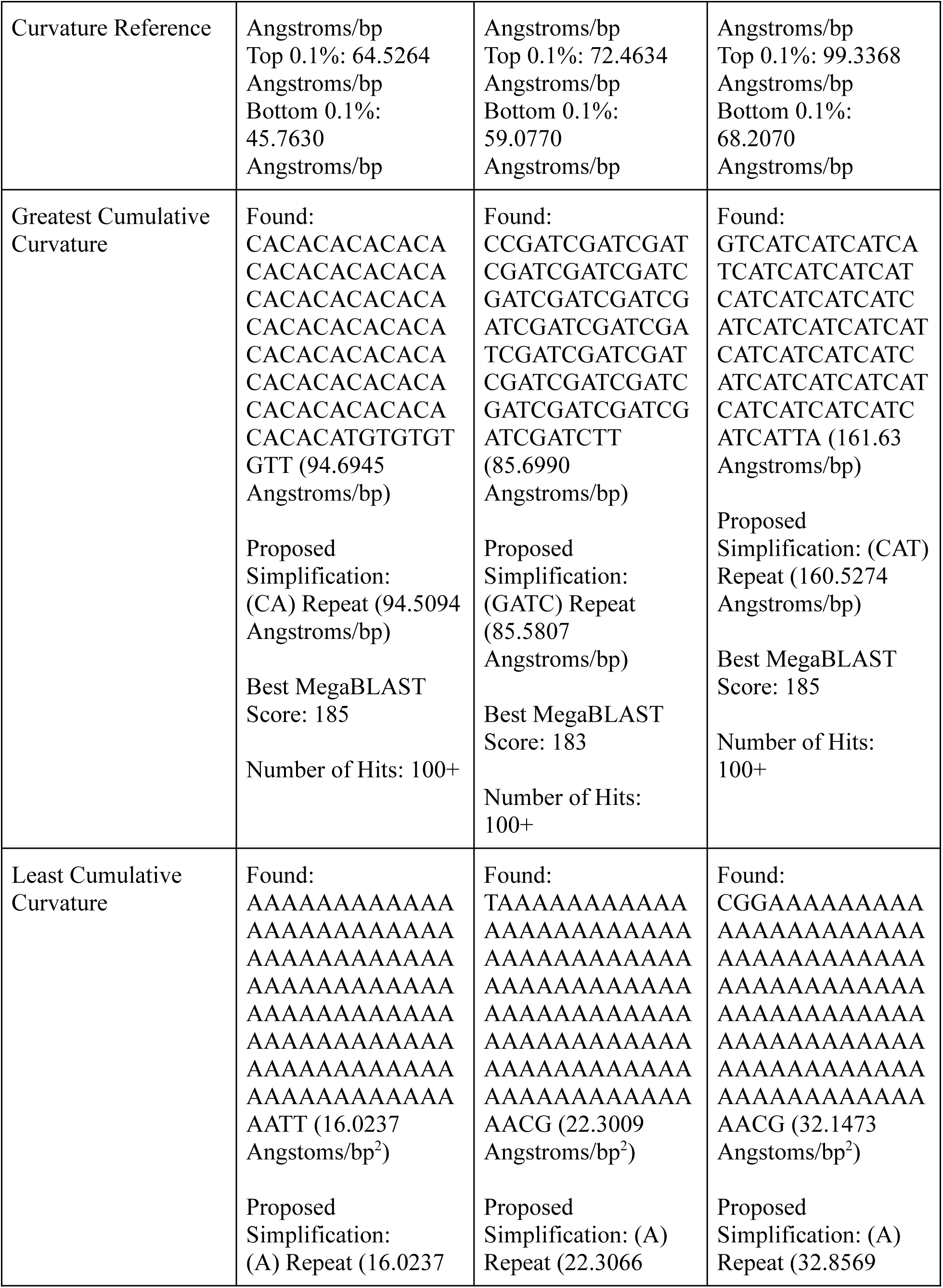
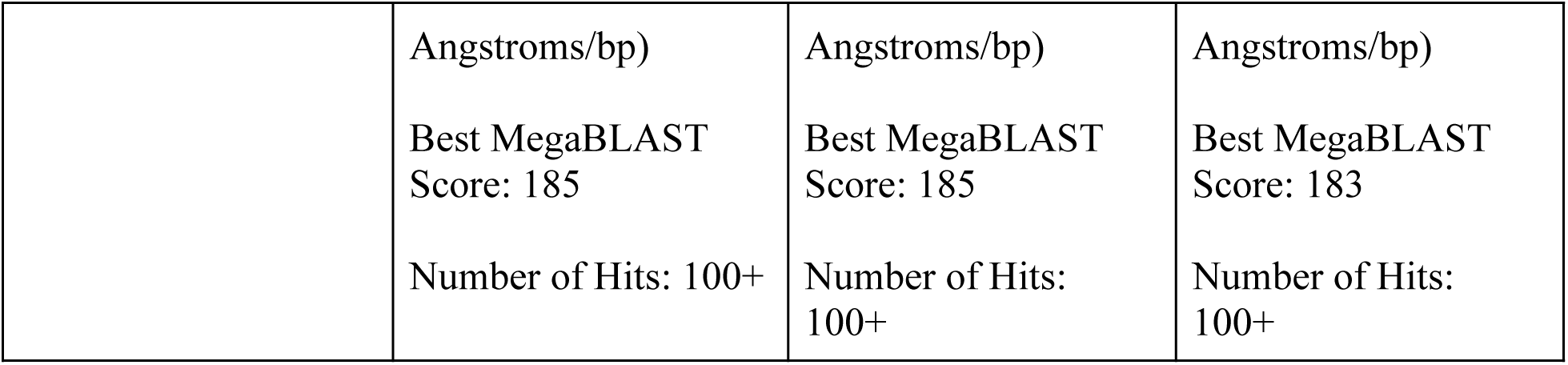
Table containing all results from searches of the DNA sequence space: Entries marked reference provide some nonparametric descriptors of the parameter’s distribution for reference. These values were obtained from 10,000 randomly generated sequences. For found sequences, first the actual sequence is given (simplified if possible) along with the value of its parameter. A simplification is given below also with parameter value if applicable. Finally, number of hits and score of top hit from a MegaBLAST of the unsimplified sequence are given.

## Proof of Periodic Extrema

### Definitions

DNA sequence can be equated to a random walk on a weighted directed complete digraph A with four vertices representing the nucleotides, with loops are allowed. By putting weights on the 16 distinct edges of the graph s={s_1_, s_2_, …, s_16_} one can express any additive function of the edges (for example the total extension) using auxiliary function f with parameter 0<r<1, where *i*(*k*) is the sequence of edges corresponding to the DNA sequence.

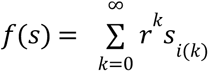

**Lemma:** Let S be the space of all infinite walks on A. Then f(s) is bounded on S.

*Proof*: There are at most 16 distinct edges for the digraph A. Thus, there exists a maximum weight edge (*s_max_*) and a minimum weight edge (*s_min_*). Thus, for any sequence s the bounds are

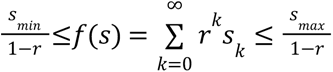

**Lemma**: f(S) contains its extrema.

*Proof*: Let X={*x*_1_, *x*_2_,…} be a sequence of points in f(S), with *x_j_* realized by the sequence *s^j^*={s^j^_i(1)_s^j^_i(2)_,…} If X is finite, there is clearly a convergent subsequence so assume X is infinite. Every element of X can be represented as

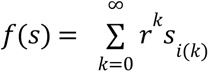

Where s is a valid infinite DNA sequence. There are at most 16 values *s_i_*_(*k*)_ could be. To find a convergent subsequence of f(S), we do the following. For every kth index, there must exist some edge weight s_k_ that occurs in an infinite number of f(s) in X, else X would not be infinite.

Starting at k=0, gradually require that the next element of our convergent subsequence of X must contain some edge weight s_k_ that shows up in an infinite number of elements of X. Limiting X to the subsequence consisting of DNA sequences where s_0_ to s_k_ are fixed, X is still infinite and we can require s_k+1_ to be fixed if it appears an infinite number of times. The subsequence will eventually converge to some f(s) where s is composed of the edges gradually fixed to produce the subsequence. This f(s) will be in f(S) and hence f(S) is sequentially compact.

Since f(S) is a subset of the real numbers and is sequentially compact, by the Bolzano-Weierstrass Theorem f(S) is compact. Since f(S) is compact it must contain both its maximum and minimum.

**Lemma**: The extrema of f(S) are eventually periodic (pre-periodic).

*Proof*: Consider the maximum of f(S). From any given vertex, the maximum must take the maximum weight walk starting at that vertex. The maximizing walk from a given vertex is independent of which index that vertex was reached at. Thus, if any maximum walk contains a cycle that cycle must repeat to be maximizing.

**Lemma**: r can be chosen such that for any r greater there is no change in extrema.

*Proof*: Since there are a finite number of vertices and edges, there are a finite number of possible cycles. Since there are a finite number of cycles of finite length, there are a finite number of subwalks of these cycles. Thus, there exists an r less than but close enough to 1 where the ordering of the distinct weights of all subwalks of all cycles on A is equivalent to when r is 1. Note that if two walks have the same weights, the extrema may not be unique.

**Table S2:** Results from performing BLAST searches on random motifs of length 11 bp repeated to 500 bp.

| Random Sequence Motif | Number of Hits with P value $<1E-50$ | Top Species | Top Bit Score |
| --- | --- | --- | --- |
| AGCTTGAATCG | 16 | *Polia nebulosa*<br>(Species of Owlet Moth) | 471 |
| TTGTCGTCCCC | 31 | *Pungitius pungitius*<br>(Ninespine Stickleback) | 617 |
| TTGGAGATAAA | 25 | *Psylliodes chrysocephala* (Species of Flea Beetle) | 489 |
| CGCGGCTCACA | 24 | *Sprattus sprattus*<br>(European Sprat) | 511 |
| TCAGTCTAGTA | 45 | *Steromphala cineraria*<br>(Species of Top Shell) | 736 |
| TATTACGGGCA | 5 | *Cyclopterus lumpus*<br>(Lumpfish) | 378 |
| CCTCCAGGGCG | 4 | *Philonthus cognatus*<br>(Species of Rove Beetle) | 360 |
| CATCTTTGTCA | 57 | *Protula* sp. h YS-2021<br>(Species of Polychaete) | 609 |
| AGAAACAAGGA | 294 | *Poecilia reticulata*<br>(Guppy) | 514 |
| GTGAAACTAAG | 17 | *Gobius niger* (Black<br>Goby) | 484 |
| TCGAATAGAGC | 6 | *Eumerus sabulonum*<br>(Species of Hover Fly) | 556 |
| CGAGTTCGATT | 119 | *Polyommatus icarus*<br>(Common Blue) | 700 |
| TCAGTTTAGAA | 28 | *Myripristis murdjan*<br>(Pinecone Soldierfish) | 506 |
| CAGTTAGGTTG | 29 | *Luidia sarsii* (Species<br>of Starfish) | 707 |
| ACCTGATCCAG | 189 | *Microchirus variegatus*<br>(Thickback Sole) | 799 |
| GAGGTAGCCGA | 11 | *Emiliana huxleyi*<br>(Species of<br>Haptophyte) | 508 |
| TTGAAACTCTT | 86 | *Blennius ocellaris*<br>(Butterfly Blenny) | 425 |
| CTAAATGAATA | 103 | *Petrosia ficiformis*<br>(Species of Sponge) | 659 |
| CGTTCGGGGTG | 17 | *Oryzias latipes*<br>(Japanese Medaka) | 399 |
| GACCCGACGAC | 50 | *Bemisia tabaci* (Species<br>of Whitefly) | 477 |

**Table S3:**
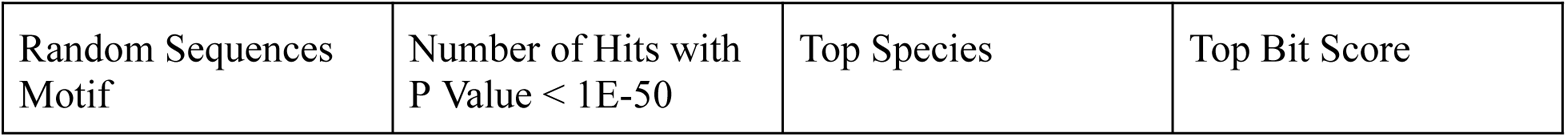
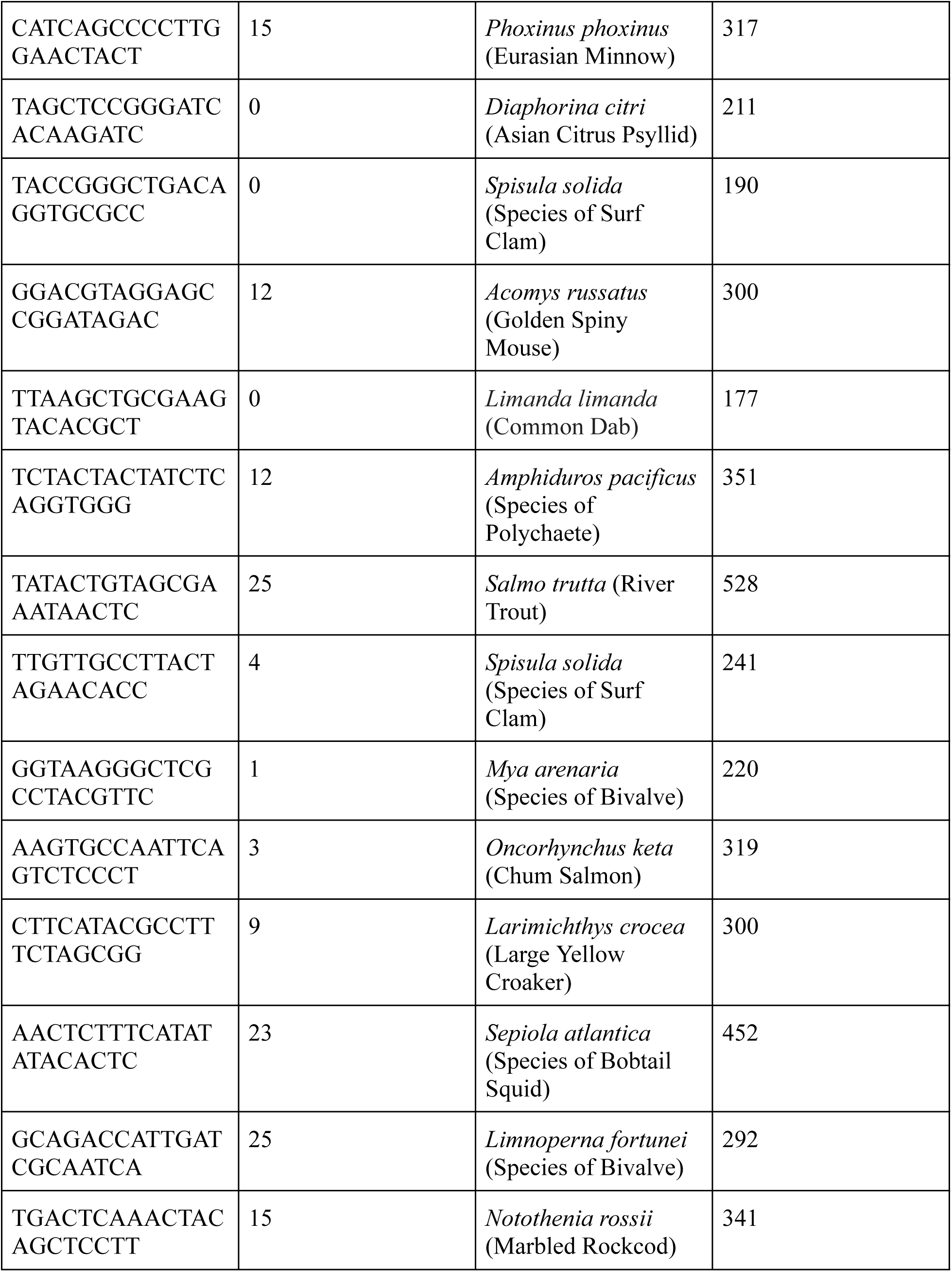
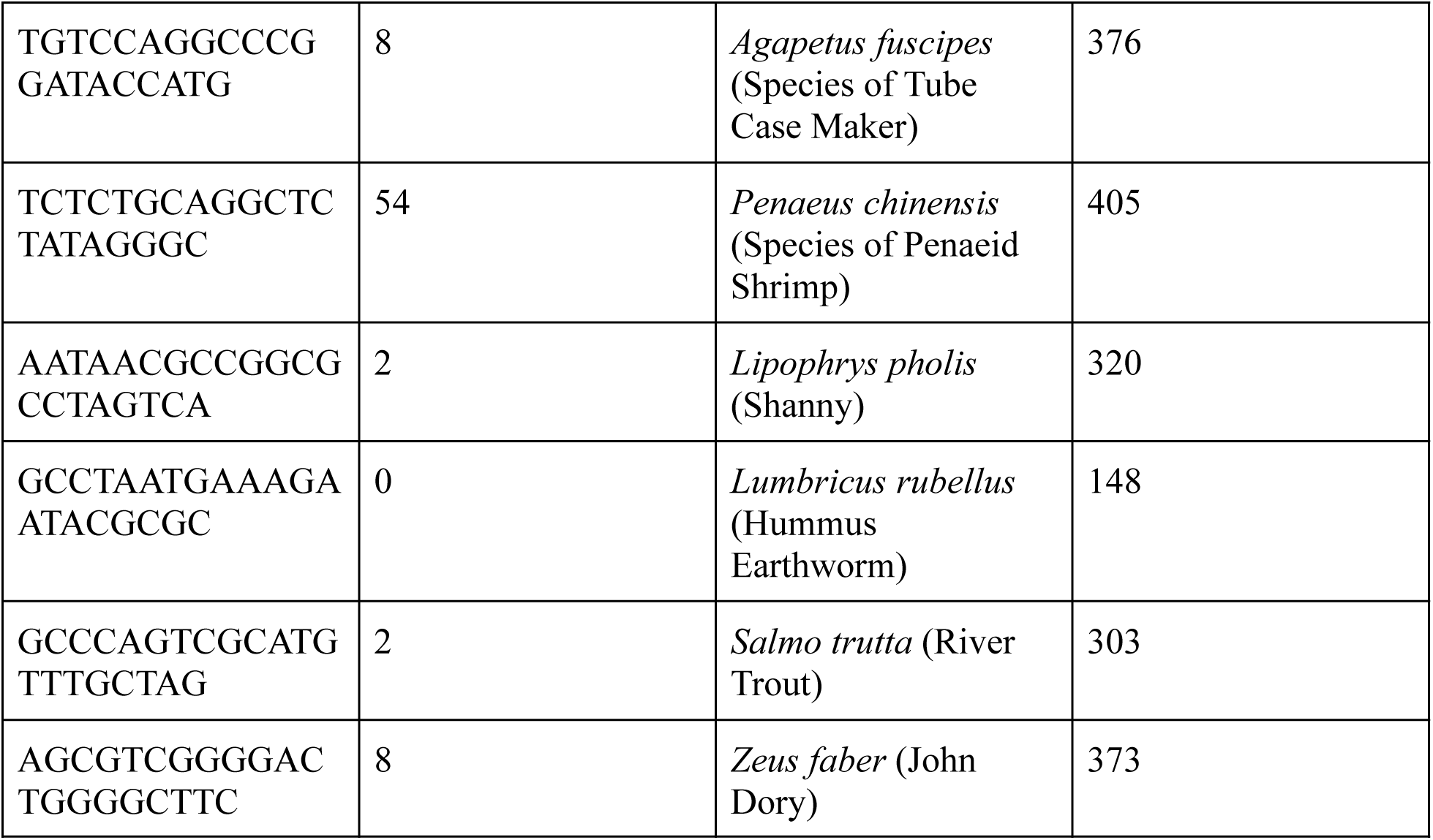
Results from performing BLAST searches on random motifs of length 21 bp repeated to 500 bp.

**Table S4:**
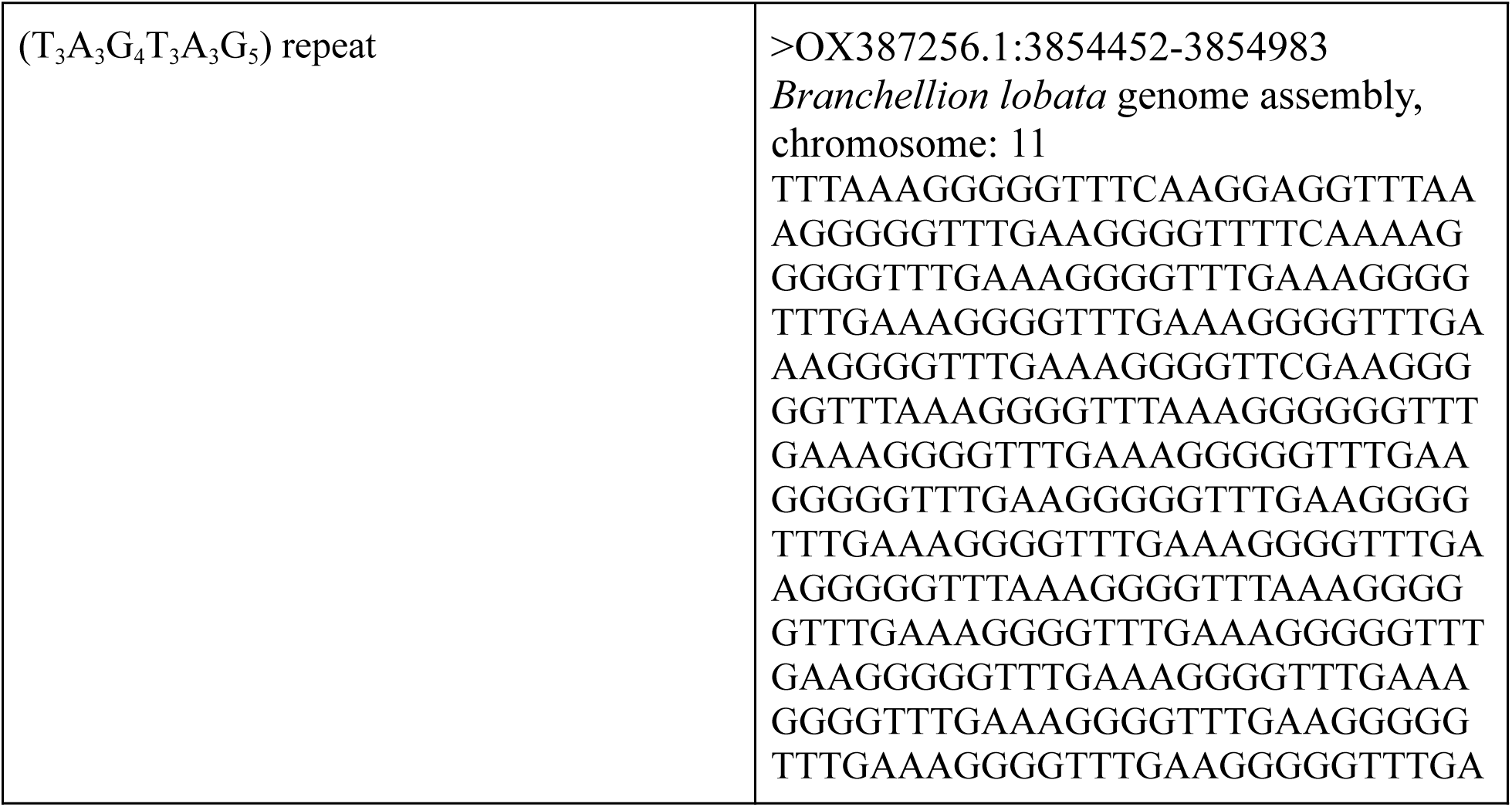
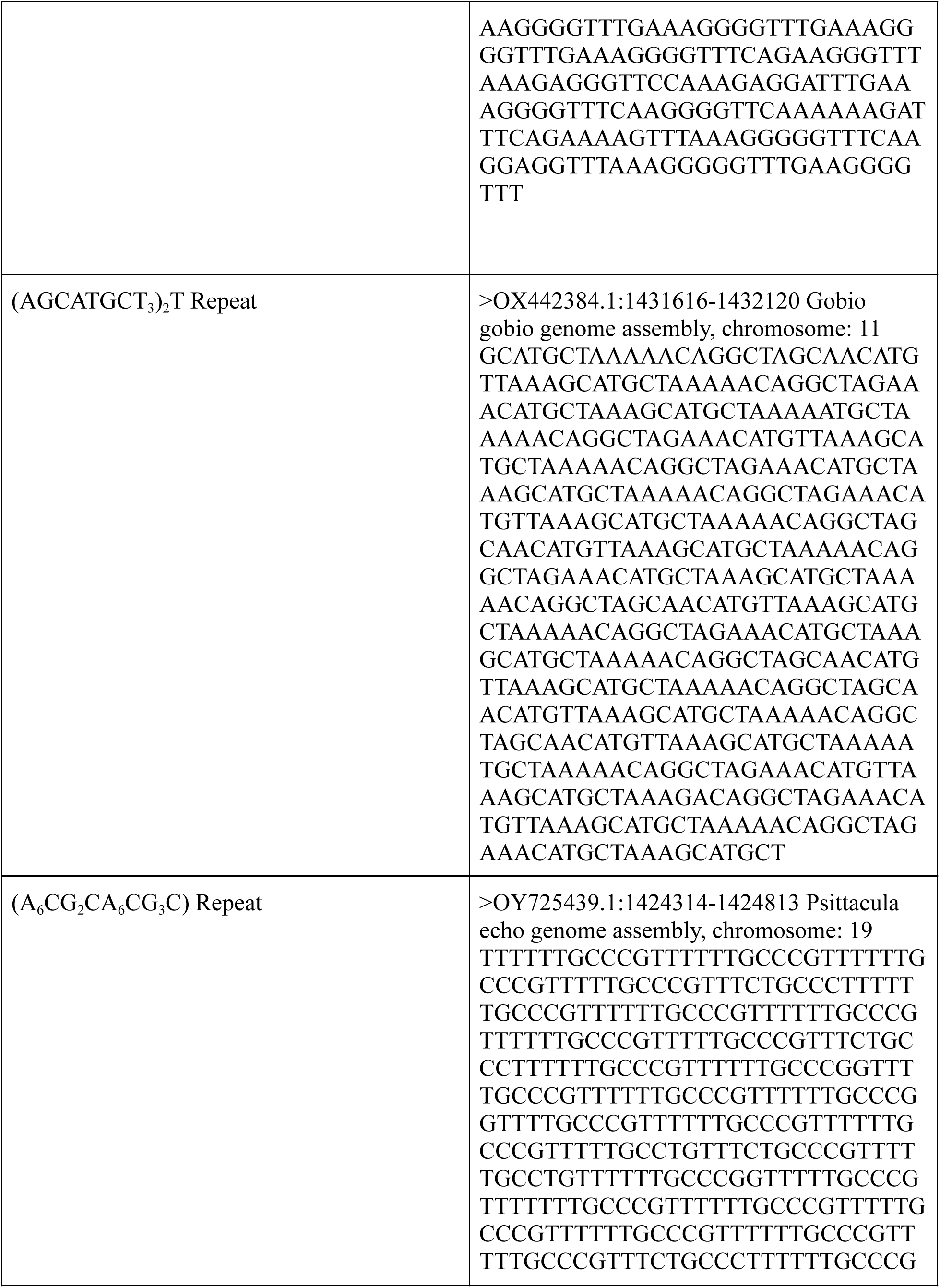
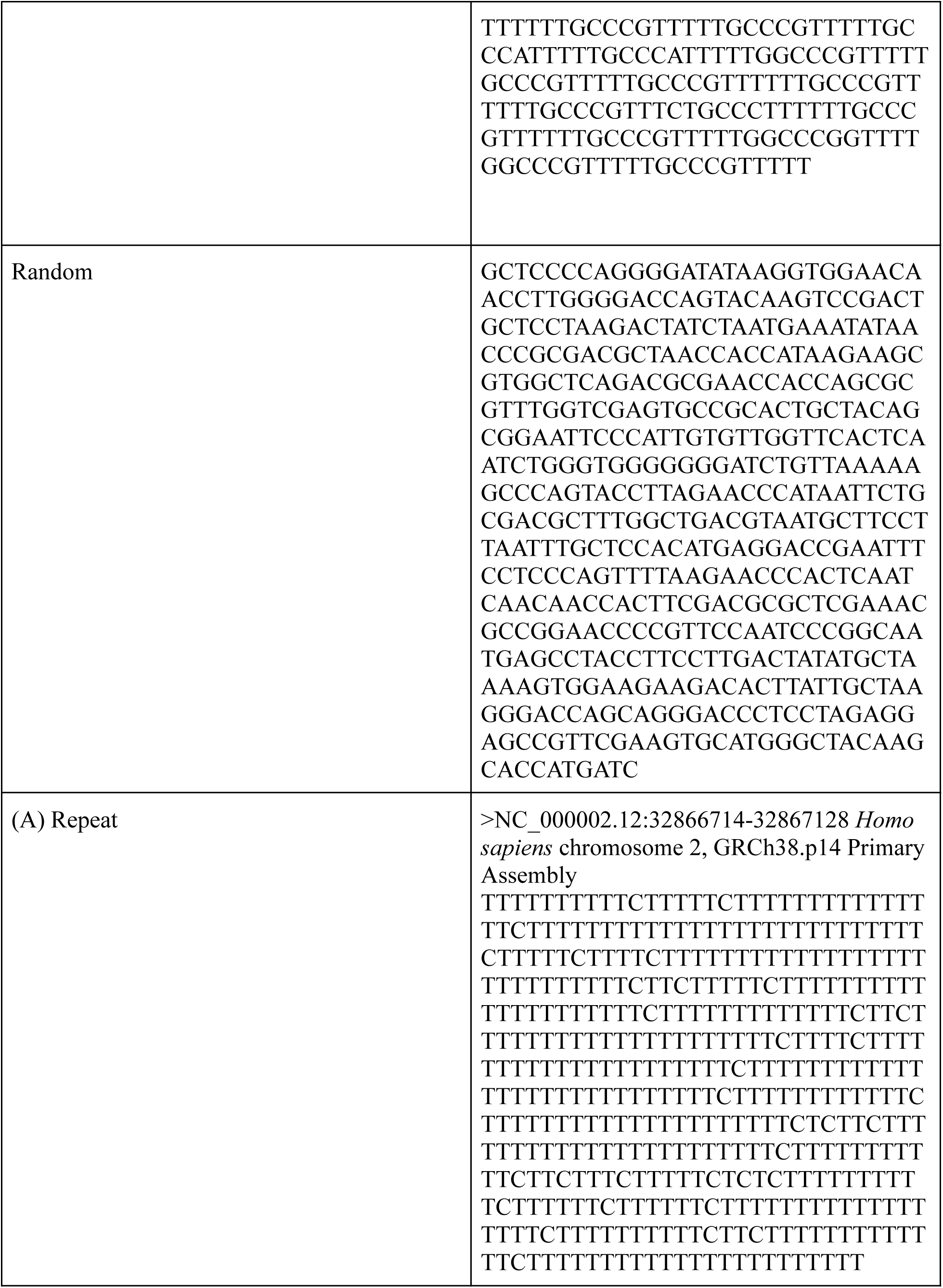
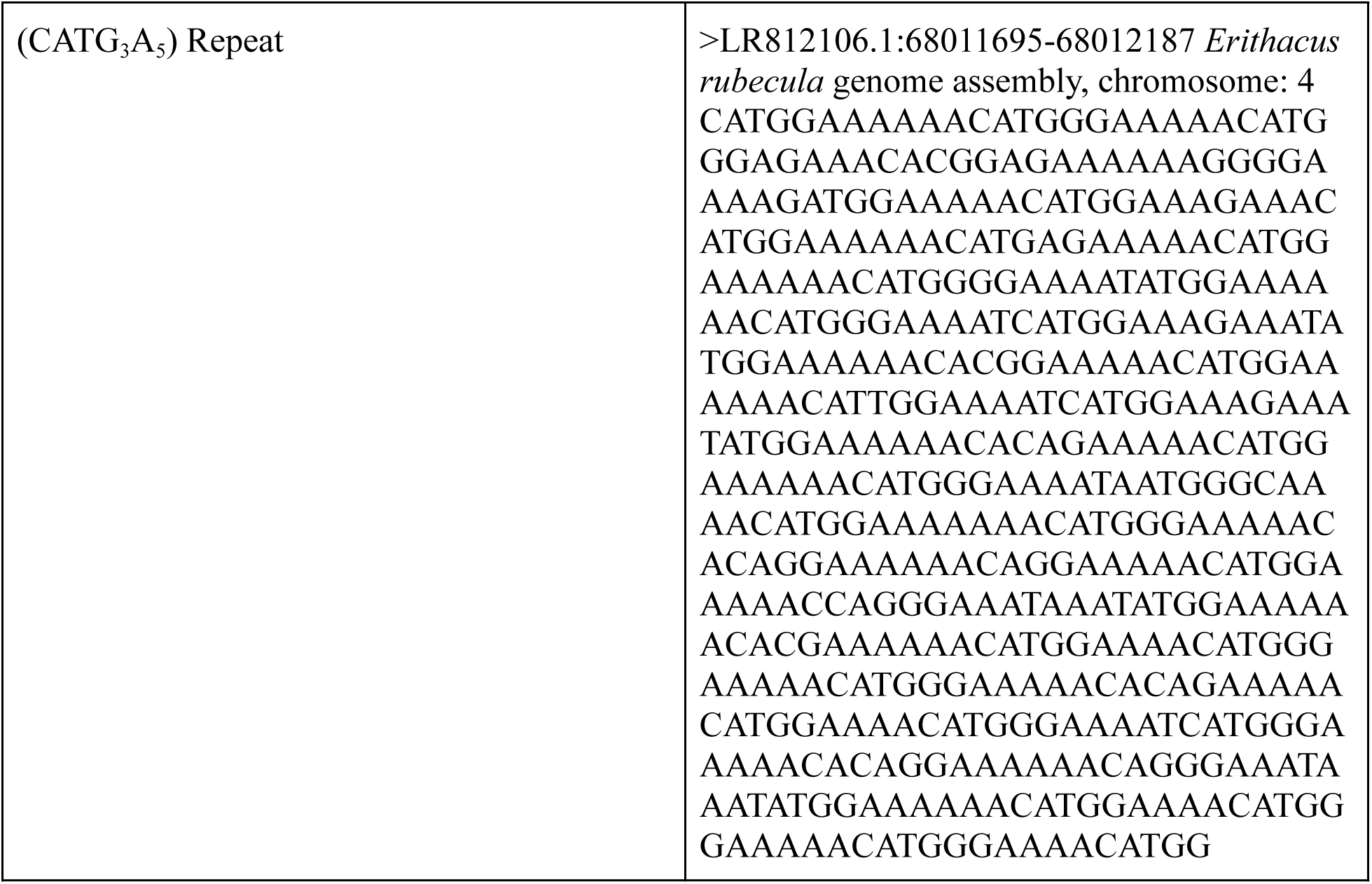
Table containing sequences used in Figure 4 along with the poly A repeat found in the human genome and the best hit from the molecular dynamics parameter search.

